# The HIV-1 ribonucleoprotein dynamically regulates its condensate behavior and drives acceleration of protease activity through membraneless granular phase separation

**DOI:** 10.1101/528638

**Authors:** Sébastien Lyonnais, S. Kashif Sadiq, Cristina Lorca-Oró, Laure Dufau, Sara Nieto-Marquez, Tuixent Escriba, Natalia Gabrielli, Xiao Tan, Mohamed Ouizougun-Oubari, Josephine Okoronkwo, Michèle Reboud-Ravaux, José Maria Gatell, Roland Marquet, Jean-Christophe Paillart, Andreas Meyerhans, Carine Tisné, Robert J. Gorelick, Gilles Mirambeau

## Abstract

A growing number of studies indicate that mRNAs and long ncRNAs can affect protein populations by assembling dynamic ribonucleoprotein (RNP) granules. These phase separated molecular ‘sponges’, stabilized by quinary (transient and weak) interactions, control proteins involved in numerous biological functions. Retroviruses such as HIV-1 form by self-assembly when their genomic RNA (gRNA) traps Gag and GagPol polyprotein precursors. Infectivity requires extracellular budding of the particle followed by maturation, an ordered processing of ~2400 Gag and ~120 GagPol by viral protease (PR). This leads to a condensed gRNA-NCp7 nucleocapsid and a CAp24-self-assembled capsid surrounding the RNP. The choreography by which all of these components dynamically interact during virus maturation is one of the missing milestones to fully depict the HIV life cycle. Here, we describe how HIV-1 has evolved a dynamic RNP granule with successive weak-strong-moderate quinary NC-gRNA networks during the sequential processing of the GagNC domain. We also reveal two palindromic RNA-binding triads on NC, KxxFxxQ and QxxFxxK, that provide quinary NC-gRNA interactions. Consequently, the nucleocapsid complex appears properly aggregated for capsid reassembly and reverse transcription, mandatory processes for viral infectivity. We show that PR is sequestered within this RNP and drives its maturation/condensation within minutes, this process being most effective at the end of budding. We anticipate such findings will stimulate further investigations of quinary interactions and emergent mechanisms in crowded environments throughout the wide and growing array of RNP granules.

## Introduction

Biomolecular condensates (BCs) are membraneless, intracellular assemblies formed by the phenomenon of liquid-liquid phase separation (LLPS) [1–5]. Several types of such assemblies have been observed inside eukaryotes with a variety of suggested functions. These range from adaptive cellular response to physiological stresses via formation of stress granules [6–9], to meeting the demands of intracellular transport or signalling, amongst many other functions [3]. They have also importantly been linked to disease [10, 11]. Fundamentally, due to their capacity to concentrate biomolecules, a suggested principal function of BCs has been that they regulate enzyme biochemistry [12–16]. Many condensates sequester mRNAs and associated RNA-binding proteins into what are termed RNA granules [17–24]. The material properties of such granules can vary depending on composition and biological functionality [25] - from dynamic architectures with liquid-like phases to non-dynamic gel-like phases [26]. Phase transitions between liquid- to gel-like phases due to condensate aging have also been observed [27].

The concept of quinary interactions [28, 29] - the emergent sum of many weak interactions that may occur in a crowded biomolecular environment - has been suggested to promote the assembly of highly stable, but dynamic and transient multi-macromolecular complexes without any requirement for membrane compartmentalisation [30–34]. Compatible with this concept, multivalent molecules that enable assembly of dense networks of weak interactions are emerging as major molecular drivers that underpin the formation of BCs [35–38]. In particular, cooperation between long polymers, such as RNAs, together with folded proteins and intriniscally disordered proteins (IDPs) may be an essential feature of many condensates [3, 39, 40]. Furthermore, constituent binding affinity, valency, liquid network connectivity and critical post-translational modifications all play a role in regulating BCs [41–48].

Recently, constituents of RNA-containing viruses, such as HIV-1 and SARS-CoV-2, have been shown to phase separate into biomolecular condensates inside cells [49], using their repertoire of IDPs [50] in conjunction with the RNA-binding capacity of their nucleocapsid proteins to interact with genomic RNA elements [51–56].

Even though an HIV-1 particle is derived from the self-assembled Pr55Gag shell and is ultimately enveloped by a lipid membrane, the concept of quinary interactions is clearly applicable in describing its dynamic assembly at the mesoscopic scale – since it forms a confined RNP gel phase in a highly crowded space, within a limited time frame and in a cooperative manner. Pr55Gag is composed from N- to C-termini of matrix (MAp17), capsid (CAp24), spacer peptide SP1, nucleocapsid (NC), spacer peptide SP2 and p6 protein. Key players here consist of NC protein intermediates with their variable nucleic acid (NA) binding properties that are dependent upon their processing state [57–60]. Tethered within the virion by approximately 2400 GagNC domains, the two single strands of 9.2 kb-long gRNA specifically scaffold Pr55Gag self-assembly. Subsequently, the HIV-1 RNP complex engages a granular condensation during the sequential proteolysis of the Pr55Gag RNA-binding domain into the mature nucleocapsid protein (NCp7) by the viral protease (PR) [59, 61, 62]. PR is derived by autoprocessing of a smaller number of GagPol within the Pr55Gag assembly that additionally contain reverse transcriptase (RT) and integrase (IN). Approximately 60 PR homodimers are potentially available to catalyze maturation, which principally requires 12000 cleavage events. Cleavage of GagNC by PR generates first NCp15 (NCp7-SP2-p6) bound to, and forming with the gRNA an RNP intermediate that physically detaches from the remaining outer MA-CA-SP1 shell. The second cleavage between SP2 and p6 releases NCp9 (NCp7-SP2) (Figure 1a). Single-stranded nucleic acids (ssNA) stimulate both cleavage events in vitro [58, 63, 64]. The third cleavage produces the mature 55 amino acids (aa)-long NCp7 and SP2. Within the virus, NCp15 seems to condense gRNA less well than NCp9 and NCp7 [65]. Yet NCp9 does not appear as functional as NCp7 [66]. NCp15 and NCp9 are short-lived species not detected during typical virus production [60]. Why such intermediates are maintained along the HIV-1 maturation process remains unclear.

**Fig 1.**
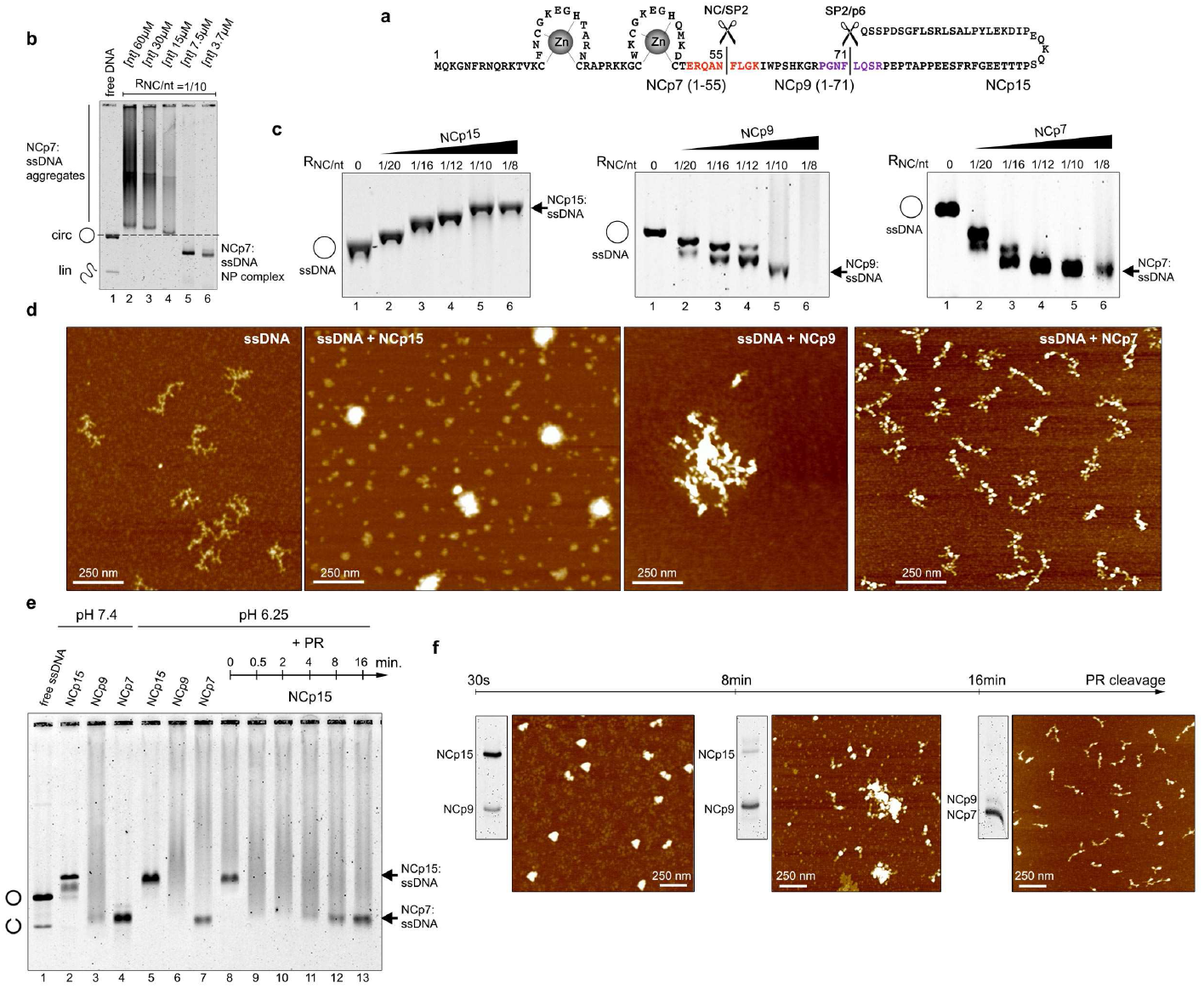
Weak-Strong-Moderate quinary properties of NCp15, NCp9 and NCp7. a, Sequence of NCp15 and PR cleavage sites. b, on agarose gel, the electrophoretic migration of M13 ssDNA:NCp7 NP complexes (1NCp7/10nt) for serial dilutions of both DNA (expressed in total [nt]) and NCp7 shows a switch from aggregation to intramolecular condensation. Position of the unbound circular (circ) ssDNA and trace of linear (lin) molecules is indicated on the left. c, M13 ssDNA:NCp15 complexes are up-shifted by EMSA upon increasing protein concentration, while ssDNA:NCp9 and ssDNA:NCp7 complexes are accelerated, thus compacted, and aggregate after NA saturation. RNC/nt indicate the protein/nucleotide ratio. d, Condensation of M13 ssDNA by NCp7 and NCp9 is visualized by AFM and show intramolecular condensates involving DNA strand bridging. NCp9 aggregate these condensates while ssDNA:NCp15 complexes appear globular. The cleavage kinetics of M13 ssDNA-bound NCp15 by PR was followed by EMSA (e), SDS-PAGE and AFM (f), showing weak-strong-moderate quinary properties of NCp15, NCp9 and NCp7, respectively, along the sequential maturation of NCp7 by PR. In the EMSA gel are shown NP complexes migration controls at pH 7.4 and pH 6.25 (optimal for PR cleavage).

HIV-1 PR is an aspartyl-protease, enzymatically active only as a homodimer. Recombinant PR is stabilized in vitro by high ionic strength (>1 M NaCl) and catalysis is strongly activated under acidic conditions (pH 5.0 or even lower). Lower salt (0.1M NaCl) and increasing the pH to 6.0 limits the acidic catalysis and shifts the equilibrium towards the monomer [67]. At quasi-neutral pH, in low salts and an excess of PR, the in vitro cleavage of Gag follows the sequential mechanism described above leading to NCp7 and the condensed RNP [68]. RNA or ssDNA promote NCp15 cleavage in vitro [58, 69], while recent reports have shown that direct RNA-PR contacts enhance the enzyme activity [64]. Consequently, PR appears to engage in an intricate partnership with NC and gRNA during viral maturation that remains incompletely understood. HIV-1 NCp7 contains a small globular domain formed with two zinc fingers (ZFs) that generate a hydrophobic pocket with two aromatic residues (Phe16 and Trp37). This platform stacks with unpaired nucleotides, preferentially guanosines exposed in RNA or ssDNA secondary structures, while basic residues stabilize the complex through electrostatic interactions with the NA backbone. With particular stem-loops in gRNA or DNA, this results in the formation of specific complexes [70, 71]. NCp7 is also a highly mobile and flexible polycationic condensing agent; like polyamines, transient protein:NA electrostatic contacts neutralize phosphate backbone repulsions lowering the overall energy of the RNP complex [59, 72, 73].

The binding properties of the various maturation states of the nucleocapsid protein to nucleic acids vary [74–76]. In vitro, these properties induce a massive co-aggregation of recombinant NCp7 and NCp9 with NA templates [57,60,77]. This quinary interaction capability guides the matchmaking/NA chaperone activity by facilitating intra- and intermolecular RNA-RNA interactions required for functional gRNA folding [78]. Such crowding effects rely on basic residues particularly concentrated in the two small flexible domains, the (1-14) N-terminal domain and the (29-35) linker between the ZFs [72]. NCp15 shows slightly different NA binding and chaperone properties but is essentially characterized by a reduced ability to aggregate NA [57, 60, 79], properties recently correlated with a direct fold-back contact between the p6 and ZF domains [60]. NCp9 shows an enhanced NA affinity due to a slower dissociation rate, as well as dramatically enhanced NA aggregating activities [57, 60, 73]. Alanine substitution of acidic residues in p6 converts NCp15 to a NA-aggregating protein similar to NCp9, while addition of a p6 peptide lowers the RNA chaperone activity of NCp7 in vitro [60]. This suggests that SP2 contains an additional NA-interaction domain, which may be masked or modulated with another NCp7 domain by intra- or intermolecular protein contacts between p6 and the NC domain.

HIV-1 maturation is mandatory for viral dissemination following sequential processes of protein and RNA self-assembly, coordinated in space and time by the enzymatic activity of viral PR [61, 62, 80]. The slow in vitro kinetics of Gag proteolysis supports a general scheme for PR to be auto-processed during the completion of budding thus driving viral maturation within free, released particles in a computed time-scale close to 30 min [81]. This model is, however, inconsistent with many observations from electron microscopy that shows i) a huge majority of free but freshly released particles in a mature form containing condensed RNP [82], ii) both capsid and budding defects in presence of PR inhibitors [83], and iii) budding and maturation defects for critical NC mutants, whereas Western blots from cell extracts detects PR-processed Gag products [82]. Such findings suggest a much closer overlap between budding and maturation than generally supposed. Importantly, suppressing both PR cleavage sites in NCp15 abolishes viral infectivity [65, 84] and results in an abnormal virion core morphology [65]. In contrast, suppression of the NCp7-SP2 cleavage site shows little effect on virus morphology and infectivity in single-cycle assays, but reverts to WT (e.g. containing NCp7) after several rounds of infection [84]. A “roadblock” mechanism affecting RT activity on a NA template has been shown to be imparted by NCp9 as well as by NCp15, which could limit large-scale viral replication, highlighting NCp7 as the optimized cofactor for accurate RNP folding and viral fitness [66].

The present study highlights how HIV-1 gRNA becomes condensed by NC proteins through the action of the RNP-sequestered PR enzyme. Reconstituted systems that model non sequence-specific binding on a large scale allowed us i) to detail the quinary effects and their variations engaged in this dynamic process as well as ii) to focus on PR action in such a quinary interaction context.

## Materials and Methods

### Proteins, Nucleic Acids and Reagents

#### Proteins

The HIV-1 NC proteins and proviral plasmids were based on the pNL4-3 sequence (GenBank accession number AF324493). Recombinant wild-type and mutants of NCp7, NCp9 and NCp15, respectively 55, 71 and 123 amino acids in length, were expressed and purified as described [60, 85–87]. The CA-SP1-NC-SP2-p6 protein expression construct was generated by PCR amplifying pNL4-3 using Gag∆MA sense primer 5’-GAT CTG GGT ACC GAG AAC CTC TAC TTC CAG ATG ATA GTG CAG AAC, NL43 OCH antisense primer 5’-GCT TGA ATT CTT ATT GTG ACG AGG GGT CGC TGC and cloning the resulting product into the homologous KpnI and EcoRI sites of pET32a (Novagen, Madison, WI). Expression construct for NCp15(1) (expressing NCp15 that can be cleaved only to NCp9) and NCp15(2) (expressing uncleavable NCp15) were generated starting with the NC-SP2- and NCp15-containing proviral plasmids of Coren et al. [84], respectively. The two constructs were generated by PCR amplifying the appropriate plasmids with NL4-3 NC sense primer 5’-CGT GGG ATC CTT AGA GAA CCT CTA CTT CCA GAT ACA GAA AGG CAA TTT TAG, NL4-3 NCp15 antisense primer 5’-GTA CGT GTC GAC TCT CTA ATT ATT GTG ACG AGG GGT CGC T and cloning into the homologous BamHI and SalI sites of pET32a. Site-directed mutagenesis of the wild-type NL4-3 NCp7 construct to generate the K3A/F6A/Q9A mutant was performed using the Agilent QuickChange Site-Directed Mutagenesis kit, with verification by NA sequence analysis, for the generation of the recombinant expression plasmid, as described [88]. The K3A mutation results from changes to nucleotides 1927 through 1929 from AAA to GCC, F6A results from nucleotides 1936 and 1937 being changed from TT to GC, and Q9A results from nucleotides 1945 and 1946 being changed from CA to GC. Proteins were expressed and purified as described [60, 85–87]. Proteins were stored lyophilized and then suspended at a concentration of 1 mg/mL in a buffer containing 20 mM HEPES pH7.5, 50 mM sodium acetate, 3 mM DTT, 20% (v/v) ethylene glycol, 200 *μ*M ZnCl2 and stored at −20°C. The concentrations were determined by measuring the UV absorbance at 280 nm using the following extinction coefficients: NCp7 and NCp7 mutants: 5690 M^−1^cm^−1^; NCp9: 11,380 M^−1^cm^−1^; NCp15: 12,660 M^−1^ cm^−1^. HIV-1 PR was expressed in E. coli Rosetta(DE3)pLysS strain (Novagen) as inclusion bodies using the expression vector pET-9 and purified as described [89, 90]. The PR domain used here bears the Q7K/L33I/L63I and C67A/C95A protective mutations to respectively minimize auto-proteolysis [67] and prevent cysteine-thiol oxidation [91]. PR was suspended, adjusted to 10-20 *μ*M concentration and stored at −80°C in 50 mM sodium acetate pH5.5, 100mM NaCl, 1 mM DTT, 0.1 mM EDTA, 10% (v/v) glycerol.

#### Nucleic Acids

The circular 7,249 nt M13 ssDNA (m13mp18) was purchased from Bayou Biolabs, the 3,569 nt MS2 RNA from Roche GmBh. Linear m13mp18 molecules were generated by annealing a complementary oligonucleotide to form a restriction site for BsrB I (NEB) as described [92]. The oligonucleotides poly d(A)13 and TAR-RNA (27 nt, 5’-CCAGAUCUGAGCCUGGGAGCUCUCUGG-3’), were purchased from Sigma-Aldrich, the short RNA fragments corresponding to individual stem-loop motifs of the Psi region: SL1 (17 nt), SL2 (23 nt), SL3 (14 nt) and SL4 (24 nt) were purchased (Microsynth) and purified by HPLC (ÄKTA design-Unicorn) on a PA-100 anion exchange column (Dionex). Plasmids used for in vitro transcription of HIV-1 RNAs used in this study have been described previously [93, 94]. Briefly, the pJCB vector was linearized with AflII, XbaI, BssHII, RsaI, or PvuII, and used as templates for the synthesis of RNAs 1-61, 1-152, 1-278, 1-311 and 1-615, respectively, by in vitro run off transcription using bacteriophage T7 RNA polymerase, followed by purification using size exclusion chromatography as described previously [95]. Likewise, plasmid pmCG67 was linearized with AvaII or SalI to produce RNAs 1-1333 and 1-4001, respectively. RNA 1-102 was obtained from a PCR product corresponding to the HIV-1 MAL sequence.

### NP complex assembly and electrophoretic mobility shift assay

Unless stated otherwise, electrophoretic mobility shift assays (EMSA) contained 1 ng/*μ*L NA templates (0.4 nM) in 20 *μ*L and were performed at 37 °C in a binding buffer containing 20 mM Tris-HCl pH 7.4, 1 mM MgCl2, 100 mM NaCl, 1 mM DTT. NCp were diluted on ice in the reaction buffer. For reactions using NCp15, the buffers were supplemented with 0.1% (w/v) Tween 20. Reactions were initiated by addition of NC as appropriate and terminated at 30 min. (unless stated otherwise) by chilling the tubes on ice and addition of 10% (v/v) a loading buffer (30% glycerol, 0.01% xylene cyanol, 10 mM Tris-HCl pH 7.4). Samples were then fractionated on 25 cm-long, 1% (w/v) agarose gels (SeaKem LE Agarose, Lonza) in 0.5x TBE. Gels were run in a Sub-Cell GT cuvette (BioRad) for 17-18 h at room temperature at 3 V/cm, stained with SybrGold (Molecular Probes) and scanned for fluorescence using a Typhoon 8600 (GE Healthcare). All experiments were performed at least in triplicate.

### Dynamic Light Scattering

Dynamic light scattering (DLS) measurements were carried out in 20 mM Tris-Acetate pH 7.4, 50 mM sodium acetate and 1 mM DTT, using 2.5 ng/*μ*L M13 ssDNA. Experiments were performed with a Zetasizer Nano-ZS (Malvern Instruments Ltd) and high precision cells (QS 3.0 mm, Hellma Analytics). Measurements were performed at 37°C, 20 min after addition of the indicated amount of NCp and analyzed by the Dispersion Technology Software provided.

### AFM imaging

NC:NA complexes were assembled under conditions used for EMSA with 1 ng/*μ*l of M13 ssDNA and in a binding solution containing 10 mM TrisAcetate pH 7.0, 50 mM sodium acetate, 2.5 to 5 mM magnesium diacetate and 0.5 mM TCEP. A freshly cleaved muscovite mica surface was pre-treated for 2 min with a fresh dilution of spermidine (50 *μ*M), extensively rinsed with water and dried under a nitrogen flow [96]. A 5 *μ*L-drop of the NP complexes was deposited on the surface and incubated for 3-5 min and dried with nitrogen gas. AFM images were carried out in air with a multimode scanning probe microscope (Bruker) operating with a Nanoscope IIIa or V controller (Bruker) and silicon AC160TS cantilevers (Olympus) using the tapping mode at their resonant frequency. The scan frequency was typically 1.0 Hz per line and the modulation amplitude was a few nanometers. A second order polynomial function was used to remove the background with the AFM software.

### Proteolysis assays

#### NC cleavage and SDS-PAGE analysis

A proteolysis assay of NCp15 bound to ssDNA using recombinant PR was showed previously [58]. The assay was optimized in this study to ensure a detailed analysis of the reaction using SDS-PAGE electrophoresis (Supplementary Fig. 3a-b). Peptides were quantified by fluorescent staining, which allowed accurate measurements in the 25-500 ng range, in agreement with our NP complex analysis. The standard proteolysis assay contained NCp proteins (6 *μ*M) incubated with NA for 5 min at 37 °C in 10 *μ*l of a PR buffer (MES 50 mM/Tris variable to adjust pH, NaCl 100 mM, DTT 2 mM, BSA 50 *μ*g/ml). Next, PR was added, unless otherwise indicated, at a concentration of 600 nM. Reactions were stopped by addition of a SDS-PAGE loading buffer and heat denaturation (5 min at 95°C), followed by 1 h incubation at 37°C in presence of 300 mM Iodoacetamide, which prevented protein oxidation (Supplementary Fig. 3a). Samples were separated on 20% acrylamide gels using Tris-Tricine SDS-PAGE in a Hoeffer MiniVE system. After migration at 160 V for 2.5 hr, the gels were fixed by 40% ethanol/10% acetic acid for 1 hr and stained overnight in 200 mL of Krypton Fluorescent gel stain (Life Technologies) diluted 1/10 in water. Gels were then rinsed with 5% acetic acid and incubated in milliQ water for 30 min before scanning with a Typhoon 8600 imager. Fluorescence counts were quantified using the ImageQuant software (GE Healthcare). Apparent Vmax was measured by dividing the product concentration by the time of incubation with [product]/ [S0] product ratio less than 30%. The PR cleavage assay of Fig. 1e-f was performed by incubating NCp15 (750 nM) and M13 ssDNA (1nM) in MES 50mM/Tris pH6.25, 100 mM NaCl, 4 mM MgCl2, 2 mM DTT for 15 min at 37°C in 50 *μ*L. PR (35 nM) was added and the cleavage was carried out at the indicated times. Each reaction was stopped by chilling the tubes on ice while a 5 *μ*l-drop was used to prepare mica for AFM, 15 *μ*l were loaded on the gel for EMSA and the remaining 30 *μ*l were used for SDS-PAGE after treating the samples as previously indicated. All experiments were performed at least in triplicate.

#### FRET assay

The proteolytic activities of PR were determined using the principles of Förster resonance energy transfer (FRET) by cleavage of a fluorogenic peptide substrate DABCYL-*γ*-abu-Ser-Gln-Asn-Tyr-Pro-Ile-Val-Gln-EDANS (Bachem, Germany) with DABCYL, 4-(4^′^-di- methylaminophenylazo)benzoyl; *γ*-abu, *γ*-aminobutyric acid; EDANS, 5-[(2-aminoethyl)amino] naphthalene-1-sulfonic acid. Incubation of PR with the probe resulted in specific cleavage at the Tyr-Pro bond and a time-dependent increase in fluorescence intensity that is linearly related to the extent of substrate hydrolysis. Kinetic experiments were carried out at 30°C in 150 *μ*L of the PR buffer (50 mM MES-Tris combination, 0.1 – 1 M NaCl, pH adjusted between 5 and 7), 5.2 *μ*M of the probe and 10-50 nM of PR. The probe was first dissolved in DMSO. The final DMSO concentration was kept at 3% (v/v). Fluorescence intensities were measured in a BMG Fluostar microplate reader. Delay time for the reaction start was calculated as the reaction slope intercept with the x axis. All experiments were performed at least in triplicate.

### Electron microscopy of HIV-1 particles

#### Maturation mutants

Mutant virions accumulating NCp15 or NCp9 were produced by transfection of mutated pNL4-3 proviral plasmids as described [84]. Plasmids were transfected into HEK 293T cells using Mirus TransIT 293 (Mirus Bio LLC, Madison, WI) according to the manufacturer’s instructions. 48 h culture supernatants were clarified and virions ultracentrifuged and examined by electron microscopy as described previously [97, 98]. At least 180 particles were analysed on the criteria that they were enveloped and a contrast was visible inside. Then the subpopulation of the diffuse cores instead of thin, dark spots was scored with, as discriminating criteria, a diameter equal or larger than 70% of the internal diameter of the particle.

#### Viral particles produced from latently infected cells

Briefly, the latently infected ACH2 cells [99] were grown under standard conditions, were plated onto 10 cm cell culture dishes at densities of 4×106 cells and incubated with or without PR inhibitor (10 *μ*M Lopinavir, Sigma). HIV production was activated by adding Vorinostat (10 *μ*M; Sigma). After 2 days, ACH2 cells were fixed with 2.5% glutaraldehyde, washed, dehydrated, embedded in epoxy resin according to standard procedures [100]. Electron microscopy images were obtained with a Tecnai Spirit microscope coupled with a 1376 × 1024 pixel CCD camera (FEI, Eindhoven, The Netherlands). We analysed 500 particles attached to the membranes after normal production and 120 after production in presence of Lopinavir (respectively 46.1 and 53% of the total number of detectable particles). Within each attached population, mainly 91% particles were identifiable, 89.6% containing a dark spot compared to only 1.4% immature for the normal population, while dark spots were not visible after viral production in presence of Lopinavir.

### Molecular Dynamics Simulations and Analysis

Molecular dynamics simulations followed a previously well-established protocol [101]. An initial structure was prepared for the NC-SP2 (RQAN-FLGK) octapeptide ligand in apo-form. Atomic coordinates for the octapeptide were extracted from the 1TSU crystal structure [101,102]. The standard AMBER forcefield (ff03) [103] was used to describe all parameters. The system was solvated using atomistic TIP3P water and then electrically neutralized with an ionic concentration of 0.15 M, resulting in a fully atomistic, explicit solvent system of approximately 14,000 atoms. Conjugate-gradient minimization was performed. The SHAKE algorithm was employed on all atoms covalently bonded to a hydrogen atom. The long range Coulomb interaction was handled using a GPU implementation of the particle mesh Ewald summation method (PME). A non-bonded cut-off distance of 9 Å was used with a switching distance of 7.5 Å. During equilibration the position of all heavy peptide atoms was restrained by a 0.5 kcal/mol/Å^2^ spring constant for all heavy protein atoms and the system evolved for 10 ns with a timestep of 4 fs. The temperature was maintained at 300 K using a Langevin thermostat with a low damping constant of 0.1/ps and the pressure maintained at 1 atm for both systems. The system was then equilibrated for 10 ns of unrestrained simulation in the canonical ensemble (NVT) with an integration timestep of 4 fs. The final coordinates were used as input for production simulations. All subsequent simulations were carried out in the NVT ensemble. All production simulations were carried out using ACEMD [104]. An ensemble of 10 × 1 *μ*s production simulations was performed. Coordinate snapshots from production simulations were generated every 10 ps, resulting in a ensemble of 106 conformers for analysis.

The octapeptide was relabelled as R52-Q53-A54-N55-F56-L57-G58-K59. The conformer ensemble was analyzed in a reaction coordinate space consisting of two order parameters: the K59-Q53 C_*α*_ distance (d_*KQ*_) and K59-F56-Q53 C_*α*_ angle (*θ*_*FQK*_). The potentials of mean force (PMF) was calculated by binning the ensemble data into microstates corresponding to the given reaction coordinate space and then calculating the mole fraction (*ρ*) of each microstate using PMF = −kBTln(*ρ*), where kB is the Boltzmann constant and T the temperature. Corresponding order parameters were calculated for each of the NMR conformers in PDB 1F6U of the NC N-terminus where d_*KQ*_ was the K3-Q10 C_*α*_ distance and the *θ*_*FQK*_ K3-F6-Q9 C_*α*_ angle. Conformers were then aligned to the NC N-terminus from PDB 1F6U by the C_*α*_ atoms of K59-R52 mapped to K3-R10 of 1F6U. The C_*α*_ RMSD was then calculated as a third order parameter (d_*n*_), its probability density was determined by binning (*ρ*(d_*n*_)) and conformers within the thresholds of 1 Å, 1.5 Å and 2 Å extracted and mapped back to the d_*KQ*_ - *θ*_*FQK*_ reaction coordinate space.

## Results

### Cleavage of NCp15 to NCp9 and NCp7 underpins Weak-Strong-Moderate quinary condensate properties

We first focused on the quinary interactions and the architectural behavior of NC:NA complexes by a combination of electrophoretic mobility shift assay in agarose gels (EMSA), atomic force microscopy (AFM) and dynamic light scattering (DLS). Examining RNPs with large ssNA templates under increasingly dilute conditions interestingly switched NCp7 binding from NA aggregation (quinary interactions) to intramolecularly-folded NP condensates (Figure 1b). NCp7 binding titrations on a circular M13 ssDNA showed a progressive process of ssDNA migration acceleration in a gel (Figure 1c), seen by AFM as tightly compact NP structures formed of folded DNA strands coated and bridged with protein (Figure 1d). Maximum ssDNA compaction was reached for saturating amounts of one NCp7 over 8-10nt [77]. Additional protein resulted in the fusion of these NP condensates into very high molecular weight structures that exhibited smearing during electrophoresis.

AFM showed a progressive accumulation of protein clusters covering the lattices while the branched and secondary structures of the ssDNA appeared melted or absent, and rather bridged into nucleofilament-like structures (Supplementary Fig.2). Omission of magnesium in the buffer (Figure 2g-i) or an excess of NCp7 resulted in fusion of the individual condensates into huge macrostructures with a spheroid shape comparable with previously described NC:NA aggregates [57, 58, 73]. NCp7 mobility was deemed necessary since this fusion was not observed and condensation was delayed at 4°C [79, 86] (Supplementary Fig 1a). The kinetics of the reaction indicated fast intramolecular condensation and a slow process of NP condensate fusion (Supplementary Fig. 1b). Low monovalent salt concentration increased NCp7/ssDNA aggregation and a strong electrostatic competition was observed with Na+ or Mg2+, as expected [73, 105] (Supplementary Fig. 1c). Mutations of key aromatic residues, Phe16 and Trp37 (Supplementary Fig. 1d-e) demonstrated that ssDNA condensation not only depend on phosphate backbone neutralization but also on base capture by the ZF domain. The apo-protein SSHS NC mutant [85] promoted DNA aggregation without acceleration of DNA mobility as expected for polycation-induced NA aggregation [106]. Finally, Ala substitution of basic residues in the N-terminal domain and the linker demonstrated these residues to be essential for ssDNA condensation, as expected. Strand circularity, e.g. the ssDNA topological constraint, favored intramolecular ssDNA bridging, whereas intermolecular ssDNA-NC-ssDNA interactions were enhanced with linear M13 ssDNA or MS2 RNA (Supplementary Fig. 1f), which demonstrated protein-NA networks involving NA-protein-NA and protein-NA-protein interactions, as proposed previously [107].

**Fig 2.**
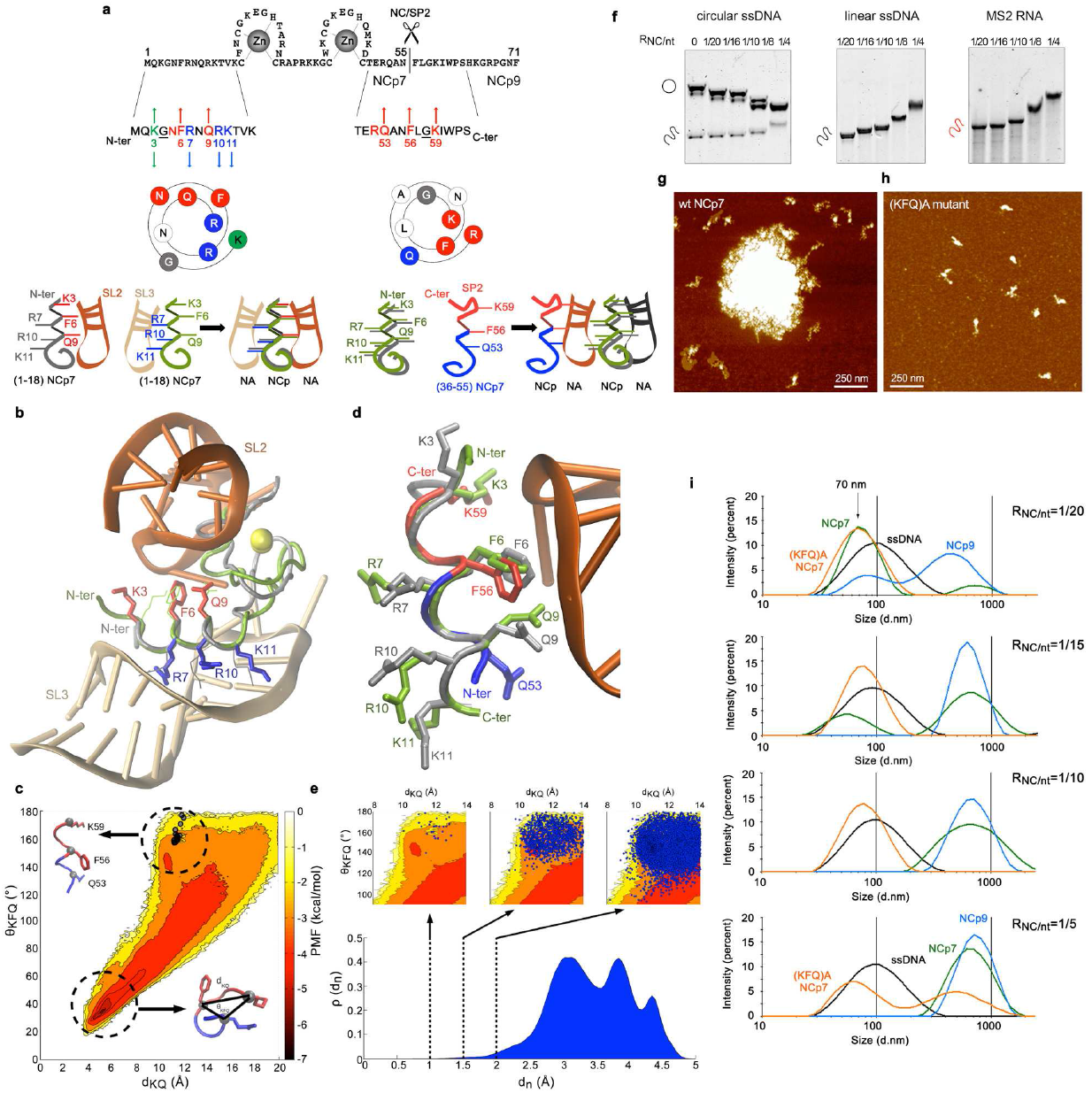
NC:NA quinary interactions. a, The N-terminal 3_10_-helix and the NC-SP2 cleavage site contain DNA strand bridging motifs. The Wenxiang diagram shows the residues of the N-terminal 3_10_-helix in contact with SL3-RNA (red) or SL2-RNA (blue). The diagram of the 53-60 sequence shows the Q53-F56-G58-K59 motif mirroring the K3-G4-F6-Q9 motif in contact with SL2. The schematics indicate possible NA-NC-NA network assemblies provided by such NA binding domains. b, Superposition of NC-SL3 (PDB ID: 1A1T) and NC-SL2 (PDB ID: 1F6U) N-terminal domain in complex with their respective NA ligands. c, The NC-SP2 apo-octapeptide exhibits substantial flexibility and is free-energetically dominated by a turn-like structure with d_*KQ*_ ~ 4-6 Å structure *θ*_*KFQ*_ ~ 20°-40° yielding a PMF of 5-7 kcal/mol. The apo-ensemble conformers sample a region with d_*KQ*_ ~10-12 Å with *θ*_*KFQ*_ ~140°-180° with less frequency (PMF~1-3 kcal/mol) where fit all NMR conformers of 1F6U, thus compatible with a 3_10_ helical structure. d, Illustration of best-fitting structural conformer of the 52-59 segment superimposed head-to-tail with the N-terminal domain of 1F6U. e, Three conformers populations were found to lie within 2 Å RMSD (9.1×10^−5^ %), 1.5 Å (1.3×10^−5^ %) and 1 Å (3×10^−7^ %) with respect to the N-terminal 3_10_ helix. Mapping the conformers onto the d_*KQ*_-*θ*_*KFQ*_ order parameters (blue circles) show they occupy the same region of the conformational sub-space.f, K3AF6AQ9A mutations in NCp7 strongly reduced the protein’s capability to aggregate circular and linear M13 ssDNA or MS2 RNA (compare with Fig. 1c, NCp7). g-h In the absence of magnesium, wt-NCp7 (1NCp7/10nt) strongly aggregates M13 ssDNA whereas the K3AF6AQ9A mutant preferentially condenses the ssDNA. i, By DLS in the absence of magnesium, M13 ssDNA (black) is condensed by the K3AF6AQ9A NCp7 mutant (orange), condensed and next aggregated by NCp7 (green) or aggregated by NCp9 (blue).

Binding of NCp9 yielded fast-migrating NP condensates for the lowest protein concentrations (Figure 1c), indistinguishable by AFM from those formed with NCp7 (Supplementary Fig. 2f). However, NCp9-driven NA condensation was seen dramatically associated with a huge fusion process by EMSA (Figure 1c), DLS (Figure 2i) and AFM (Figure 1d, Supplementary Fig. 2f-g). Linearity of the ssDNA template resulted in a huge aggregation, demonstrating the presence of an additional NA binding site in SP2 reinforcing NA-NC-NA networks (Figure 2). In contrast to NCp9 or NCp7, reaching a plateau of one NCp15 per 10-12nt, NCp15 binding to any of the three templates yielded NP complexes of lower gel mobility upon protein addition (Figure 1c), similar to canonical ssDNA binding proteins [96]. AFM visualization showed passive ssDNA coating instead of bridging compaction within individual complexes for limiting NCp15 concentration, which then led to globular structures at saturation (Fig.1d, Supplementary Fig. 2h-i). NCp15 and NCp7 retain equivalent net charges [66] (NCp15 pI 9.93; NCp7 pI 9.59). Therefore, NCp15 binding does not actively compact and aggregate ssNA, confirming previous results [57, 60]. With free NCp15, p6 has been proposed to bind to the NC domain [60]. Like NCp15 from HTLV-1 [108], HIV-1 NCp15 binding to NA might invoke quinary intermolecular p6-NC contacts instead of quinary NA-NC contacts. These interactions may freeze these globular structures and mask or block the NC residues responsible for NA compaction/aggregation. Followed by EMSA, SDS-PAGE and AFM, the dynamics of quinary NC-NA interactions through cleavage of M13 ssDNA-bound NCp15 is verified (Figure 1e-f) from weak, in the presence of NCp15, to strong with NCp9, and moderate with NCp7. From globular (NCp15), individual NP complexes are progressively converted into intramolecular condensates (NCp7) after an intermediate step of fusion (NCp9).

### Transiently unmasked NC binding sites enable modulation of NC:NA molecular interactions

A superposition of the N-terminal 3_10_ helix from the NMR structures of NCp7-SL2 and NCp7-SL3 complexes is shown in Figure 2a-b and reveals two slightly different NA backbone binding motifs for this domain, which could be virtually sandwiched between two RNA stems, providing a bridge to form RNA-NC-RNA networks [107]. Three additional basic residues over the sixteen present in SP2 poorly explain the dramatic enhancement of the NA quinary capabilities of NCp9. Examination of the NCp9 primary sequence reveals that the NC-SP2 cleavage site surprisingly contains 5 of the 8 residues of the NCp7 SL2-binding motif Lys-Gly-x-Phe-x-x-Gln-Arg, but oriented in reverse, from C- to N-terminus (Figure 2a,d).

To determine if conformers of this sequence would be compatible with a NA binding site structurally similar to those of the N-terminal 3_10_-helix, we performed all-atom molecular dynamics (MD) simulations of an NC-SP2 octapeptide cleavage site. Results of the simulations showed a large conformational area corresponding to a predominantly disordered peptide, similar to other disordered peptide regions in HIV-1 [109,110]. However, three conformer populations were found to lie within 2 Å to 1 Å RMSD with respect to the N-terminal 3_10_-helix (Figure 2c-e), as expected. The K3A/F6A/Q9A-mutation in NCp7 mostly abrogated ssNA aggregation but maintained ssDNA M13 condensation, suggesting this triad to be mostly involved in quinary interactions stabilizing NA:NC networks (Figure 2f-h). A DLS analysis in low magnesium finally demonstrated a NCp7/NCp9-driven compaction of M13 ssDNA from 100 nm to 70 nm, followed by a massive fusion/aggregation of these complexes (Figure 2i). In contrast, the K3A/F6A/Q9A NCp7 mutant was strongly defective in the fusion/aggregation process. Altogether, these data strongly support a model where the Lys(3/59)-Gly(4/58)-x-Phe(6/56)-x-x Gln(9/53)-Arg(10/52) octad would act in both NCp7 and NCp9 as a quinary interaction module, establishing bridges between NC-NA complexes at NA saturation (Fig.6a). These positions are highly conserved amongst all the HIV-1 subtypes, except at position 3 where the conservative K and R residues are found equiprobable.

### Quinary cooperation between NC and RNA drives PR sequestration and RNA-length-dependent catalytic acceleration

Followed by SDS-PAGE under conditions optimized for peptide quantification (Supplementary Fig. 3a-b), in vitro processing of the C- and N-terminal extremities of the NC domain in an environment unfavorable for PR dimers (0.1 M NaCl, pH 6.25) reveals a dramatic acceleration of NCp15, NCp9 and NCp7 production in the presence of ssNA templates (Figure 3a, Supplementary Fig. 3c-e). 100% of ssDNA- or RNA-bound NCp15 were cleaved in two distinct steps producing NCp9 and then NCp7 within minutes, confirming a distributive reaction without consecutive cuts upon the same NCp15 copy (Supplementary Fig. 3c). Without NA, complete NCp15 cleavage occurred but at a slower rate, only under acidic (pH 5.0) and high salt (1.5M NaCl) conditions (Supplementary Fig. 3d), also concomitantly producing a shorter product (NCp7*). Similar effects were observed with NA for NCp9 cleavage (Supplementary Fig. 3e) and the NC-SP2 cleavage appeared 2-3 times slower than that of SP2-p6 either starting from NCp15 or NCp9, but was completed in minutes, much faster than previously shown [68]. MS2 RNA activation also occurs for the SP1-NC site of a GagΔMA protein (CA-SP1-NC-SP2-p6), confirming previous results using a GagΔp6 (MA-CA-SP1-NC-SP2) protein [63] (Supplementary Fig. 3f).

**Fig 3.**
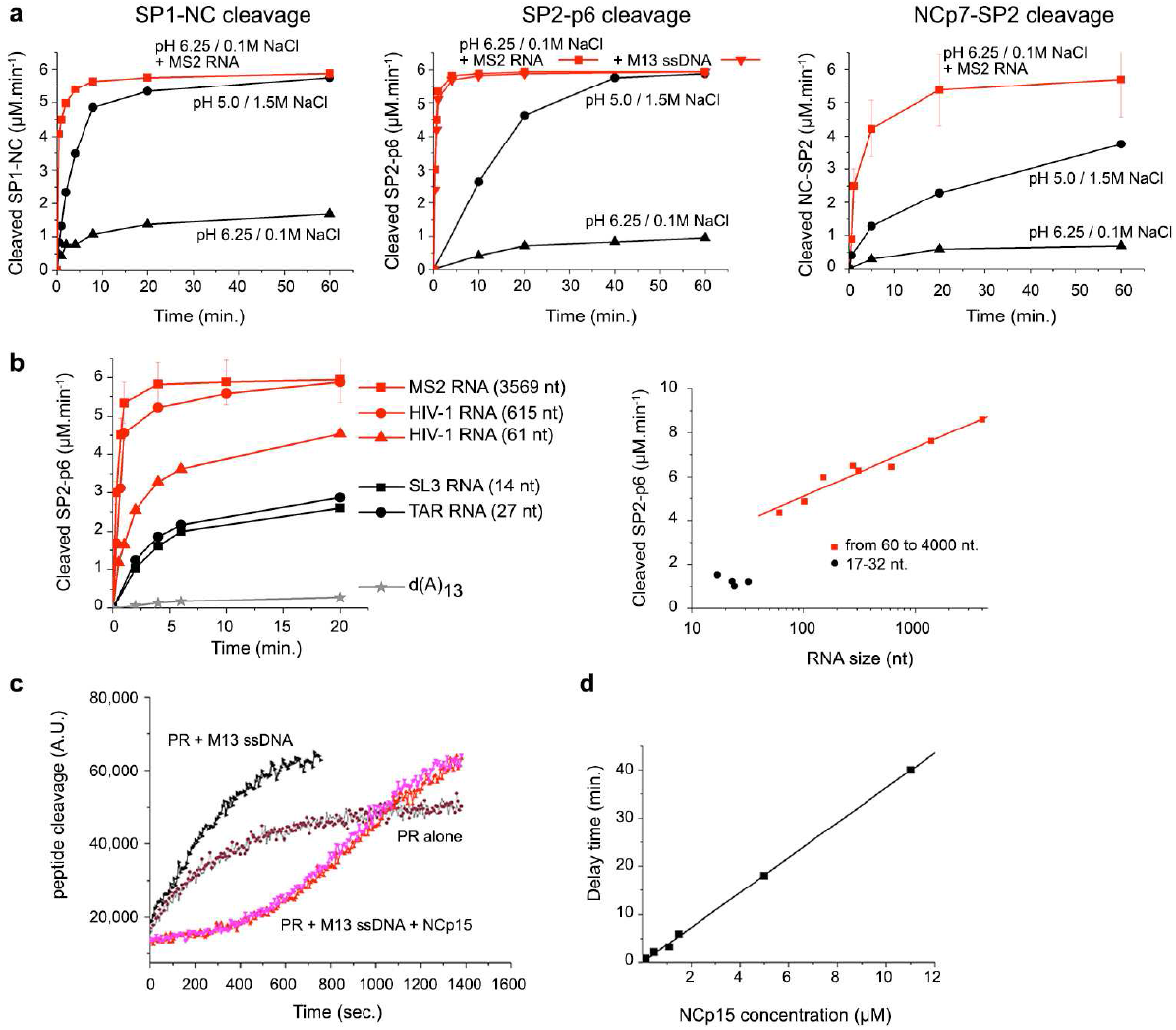
Nucleocapsid maturation is promoted by NA through sequestration of PR. a, NC binding to M13 ssDNA or MS2 RNA strongly activates in vitro cleavage of SP1-NC, SP2-p6 and NCp7-SP2 sites by PR under unfavourable conditions (pH 6.25 and 0.1 M NaCl; see Supplementary Fig. 3a-b for assay details). b, In vitro cleavage of NCp15 by PR is activated in a ssNA length-dependent manner. c, Cleavage of a MA-CA probe by PR is activated by ssDNA and is delayed in presence of ssDNA:NCp15 complexes as the latter sequester PR. MA-CA proteolysis was followed by FRET. d, Delay of the FRET probe cleavage extrapolated from (c) as a function of M13 ssDNA:NCp15 complex concentration (RNCp15/nt =1/20).

We focused on the NC-ssNA NP assemblies and their influence on NCp15 cleavage at pH 6.25 and 0.1 M NaCl. We first compared the influence of large-scale assembly of NCp15 on M13 ssDNA or MS2 RNA versus stoichiometric complexes formed between NC and a TAR RNA stem-loop (Supplementary Fig. 4a). The NA concentrations were varied for a fixed concentration of NCp15. A biphasic effect was observed in presence of either long ssNA, PR reaching maximal activity when NCp15 saturated the ssNA lattices (Supplementary Fig. 4b). A substantial reduction in PR efficacy was observed upon dispersion of NCp15 over the lattice, even though cleavage was maintained at a much higher level than in absence of NA. In contrast, neither a biphasic effect nor a rapid rate was observed in presence of the TAR-RNA.

This NA chain-length effect was next followed for NCp15 cleavage, maintaining equal nt concentration for various HIV-1 RNA stem-loops and fragments from 61 to 615 nt (Figure 3b, Supplementary Fig. 4c). A weak NC substrate, d(A)13 oligonucleotide, was ineffective in stimulating PR activity, whereas the TAR and SL3 RNA led to significant, but incomplete stimulation of the reaction. Above a critical threshold (~50 nt-length) and for a pH optimum around 6.3 (Supplementary Fig. 4e-f), PR activity scaled non-linearly with RNA length irrespective of biological origin (Figure 3b). Without NA, a 5-minute incubation between NCp15 and PR in 0.1 M NaCl at pH 6.25 resulted in no cleavage, while the addition of MS2 RNA immediately boosted the reaction (Supplementary Fig. 4d). Diluting PR for fixed NC:PR (10:1) and NA:NC (20nt.:1) ratios revealed a process resistant to dilution for NA larger than 615nt (T1/2 from 0.15 to 1 minute), whereas a strong rate decrease was observed for TAR or cTAR structures (T1/2 extrapolated to 3 h when considering the first quarter of the reaction; (Supplementary Fig. 4e). A regular decrease at pH 5.0 and 1.5 M NaCl was observed in absence of NA (T1/2 from 4 to 25 min).

These large NA chains greatly stimulated NCp15 cleavage at 0.1 M salt, with a remarkable pH optimum between 6.0 and 6.5 (Supplementary Fig. 4f). The NCp7* extra-cleavage, previously described, corresponds to a site at position 49-50 as a result of ZF destabilization at low pH [111]. This product was examined in two NCp15 cleavage mutants (Supplementary Fig. 4g) and was due to the cumulative effects of both MS2 RNA and acidic pH, which clearly “overcut” the NC domain at pH 5.4. A mildly acidic pH appears therefore beneficial in reducing irregular cleavages of NCp7 upon high PR turnover. With such optimized conditions that satisfy both PR efficiency and NC folding while restricting NCp7 cryptic site cleavages, we confirmed the RNA length effect for the NCp9-to-NCp7 reaction (Supplementary Fig. 4h-i), leading to a maximal observed rate close to the T1/2-value of NCp15-to-NCp9 reaction rate, under conditions where NCp9 and NA appeared strongly aggregated (see Figure 1).

Finally, in presence of 1:1 NC ligands (TAR, cTAR, SL3), bound NCp15 appears almost individually distributed in the reaction mix and allow a reaction-diffusion mechanism that accelerate PR turnover but under conditions where native PR is much less stable. Any substance able to increase the local concentration of either the substrate or the enzyme, or both, drives the reaction in the forward direction enhancing enzyme turnover. As such, ssNA length-dependent activation engages a NC crowding effect with NCp15 molecules coating the NA lattice and forming clusters trapping PR independently of the NP complex concentration. These NA-scaffolded clusters allow faster PR turnover, making both SP2-p6 and NC-SP2 cleavages much more efficient. In other words, the NA quinary capabilities of the NC domain induce an RNA-driven sequestration effect on PR.

To better understand this sequestration phenomenon, we used a FRET-based assay that measures the cleavage rate of a MA-CA octapeptide probe in presence of NA and/or NCp15. The assay firstly confirmed the reduction of PR activity upon pH increase and salt dilution (Supplementary Fig. 5a-b). The M13 ssDNA appeared as an effective substitute to high salt and boosted PR activity by a factor of 10 at pH 5.0, three times greater than in presence of 1.5 M NaCl. A similar effect also occurred at pH 6.25, although PR activity was strongly attenuated. These results confirmed that non-specific PR-NA interactions result in enzyme activation [64]. Adding an equimolar amount of NCp15 at pH 5.5 did not affect the reaction. In contrast, addition of NCp15 bound to ssDNA resulted in a total inhibition of the octapeptide cleavage for ~5 min, before reaching a velocity analogous to that measured in presence of ssDNA alone (Figure 3d, Supplementary Fig. 5c). PR is thus sequestered into the NP complex and completes NCp15 processing prior to cleaving the MA-CA peptide at a rate comparable with ssDNA alone, the delay time being directly proportional to NCp15 concentration with a fixed NCp15:ssDNA ratio (Figure 3d).

These data were interpreted by devising a two-substrate model of NA-modulated enzyme kinetics (Supplementary Note), which partitions the reaction volume between distinct regions that are either occupied (pervaded) or unoccupied (un-pervaded) by NA (Figure 4a). Reacting species (S1: MA-CA, S2: NCp15, E: PR) exhibit equilibrium absorption (K_*S*1_, K_*S*2_ and K_*E*_) between these regions due to nonsequence-specific NA-binding. In our model, the enzyme escape rate depends on RNP contiguity - the contiguous number (c = [NCp15]n_*l*_/[n_*t*_]) of NCp15 molecules bound per NA and not just chain length or NC-NA loading ratio alone. The phase transition is consistent with a minimum critical contiguity threshold (c_*crit*_) required to alter enzyme escape rate. Contiguity is still length dependent: fitting a reduced single-substrate model onto the experimental data (Figure 4b) yields non-linear dependence of K_*E*_ on contiguity with exponent ξ ~0.4. and c_*crit*_ ~3. By expanding to a two-substrate competitive assay (Supplementary Note) and incorporating the effects on differential enzyme decay (Figure 4d), our model is fitted to, and is compatible with the observed sequestration effect (Figure 3c) giving ξ ~1.2 and c_*crit*_ ~3. Sequestration is thus due to reduction in the RNP contiguity across the time-course of the reaction – initially the enzyme is absorbed into the RNP (K_*E*_ ≫ 1), after significant processing it is expelled (K_*E*_ ≪ 1). The model follows NCp15 cleavage directly (Figure 4c) and when scaled to in virio concentrations of enzyme and substrate as well as increased NA length, predicts a core condensation time of ~5 min (Figure 4e). Our model shows that local crowding within the RNP induces cumulative non-linear effects on non-specific enzyme binding. The absorption equilibrium constant itself depends on this local environment, consistent with quinary interactions between PR, RNA and NCp15 [30].

**Fig 4.**
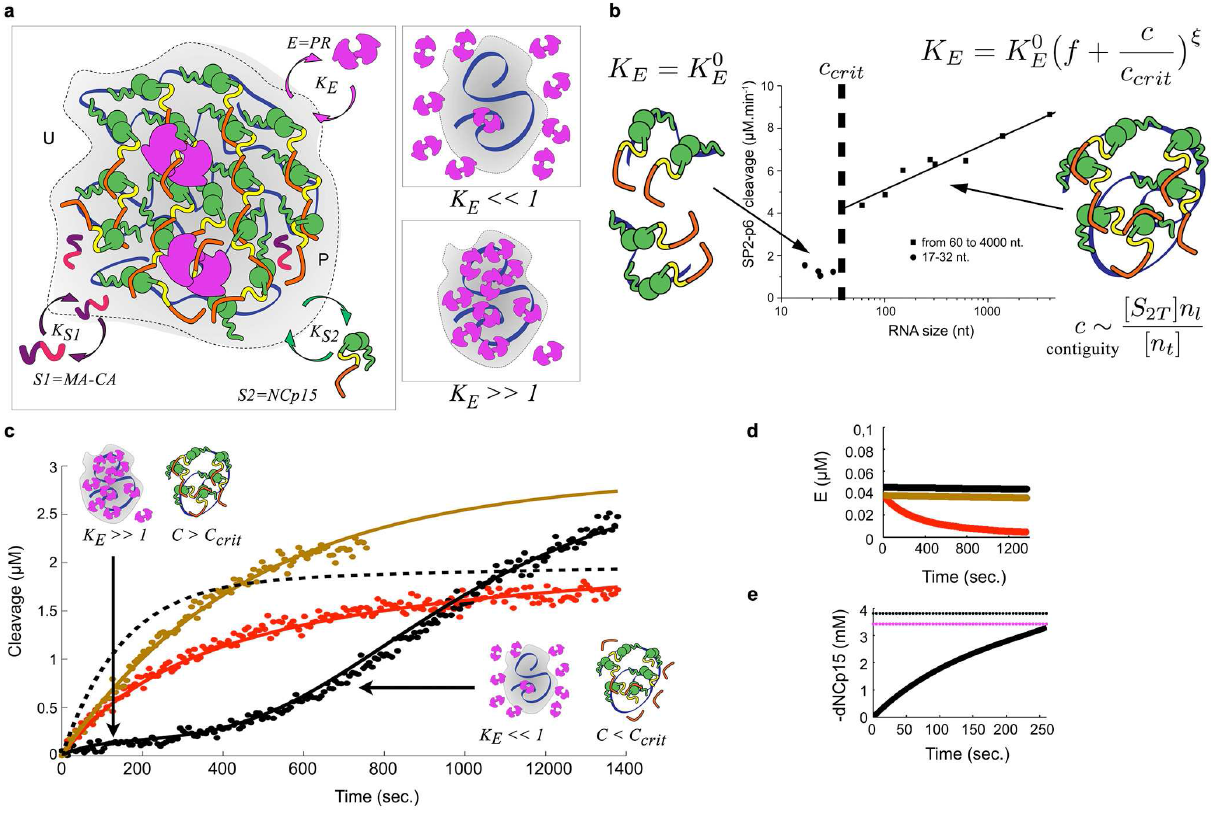
Kinetic model of two-substrate (S1, S2) processing by an enzyme (E) in a RNP. a, Reaction rate is governed by a combination of effective concentration in a volume domain and absorption kinetics for different species. For K_*E*_ ≫ 1, PR is sequestered into the RNP-pervaded volume whilst for K_*E*_ ≪ 1 it is forced out. b, A one-substrate model is fit to experimental data to determine non-linear (exponent ξ) K_*E*_ dependence on the contiguous number c of S2 molecules bound per NA above a critical threshold, ccrit. c, Fitted two-substrate model of sequestration (black) with competitive substrate alone (red) NA-present competitive substrate (brown) data. The early reaction is dominated by high contiguity (c > c_*crit*_, K_*E*_ ≫ 1) inducing enzyme sequestration. This effect dissipates upon processing (c_*crit*_, K_*E*_ ≪ 1). 90% of NCp15 cleavage (dashed line) occurs within 400 s. d, Differential enzyme decay: NA stabilizes the PR dimer; the NA-absent reaction decays with experimental half-life. e, Scaling to in virio conditions yields 90% NCp15 processing (purple) within 260 s.

### Condensate-driven accelerated PR processing temporally couples budding to maturation

In order to approach this process of RNP condensation in virio, we finally compared by TEM the core content of HIV-1 NL4-3 virus particles assembled with Pr55Gag containing un-cleavable NC-SP2 or NC-SP2-p6 sites, thus accumulating NCp9 and NCp15 respectively [84] (Figure 5a and Supplementary Fig. 6a).

**Fig 5.**
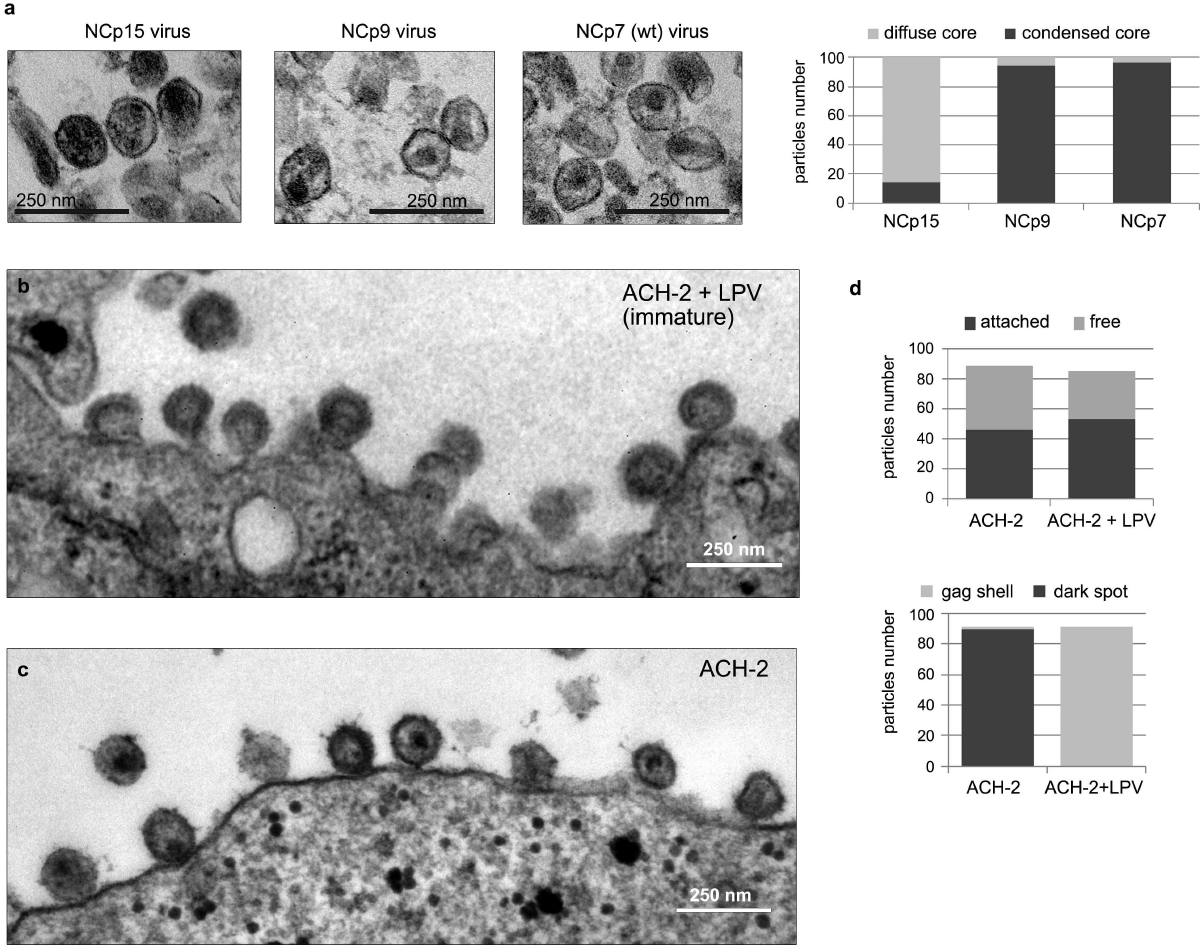
Nucleocapsid condensation within HIV-1 particles depends on NCp15 processing and is detectable in membrane-attached particles. a, By TEM, HIV-1NL4-3 virions accumulating NCp15 present defects in nucleocapsid condensation while NCp9- and NCp7-containing viruses show correct core condensation into electron-dense dark spot. Quantitation was done for close to 180 counted particles. b-c, The majority of membrane-attached HIV-1 particles produced by latently infected ACH-2 cells are immature particles in presence of LPV (b), they contain an electron-dense dark spot indicative of nucleocapsid condensation without LPV (c). Quantitation was done for close to 200 particles for LPV-treated ACH2 cells and to 1000 particles for non-treated ACH2 cells (d).

More than 90% of both NCp9- and NCp7-containing viruses display a morphologically well-formed conical capsid encasing an electron-dark spot corresponding to a condensed RNP. In contrast, more than 80% of the NCp15-containing viruses display electron-dark diffuse cores. This demonstrates that the strong-quinary NCp9 intermediate actively triggers nucleocapsid condensation, thus reducing the occupied volume and facilitating capsid rearrangement. We next imaged plasma membrane-attached particles of HIV-1 virus produced from latently-infected ACH2 cells. Washing the cell suspension before fixation enriched the proportion of attached particles engaged in budding. In the presence of a PR inhibitor, all membrane-attached particles appeared immature with a typical electron-dense Gag shell and a bottleneck that characterized budding intermediates (Figure 5b,d). Without inhibitor, most of the attached particles exhibited a dark spot and a closed envelope (Figure 5c-d). Therefore, the maturation step involving strong-quinary NCp9 occurs visibly in a time frame consistent with both the end of budding [112–114] and our kinetic model: budding and maturation appear temporally coupled.

## Discussion and Conclusions

We describe in this study HIV-1 nucleocapsid maturation as a dynamic RNA granule processing phenomenon, involving differential RNA binding activities of the NC domain that are dependent on processing state. Weak NC-RNA contacts fit with the concept of quinary interactions [28] that lead to gRNA condensation in the context of an RNA-directed gel phase [25]. We propose that this RNP follows a dynamic weak-strong-moderate (WSM) quinary model resulting in granular phase separated RNP condensation (Figure 6) with a distributive three-step processing mechanism in the order of SP1-NC, SP2-p6 and NC-SP2. Each step alters NC-RNA interaction strength within the confined gel phase. The variations in condensing the RNA (in vitro condensation plus aggregation) appear therefore directly linked with the number of amino acid residues weakly contacting NA chains, these contacts being severely limited in NCp15 due to p6 interfering with NC-SP2 NA binding [60, 66] and/or competing with the NA for binding to the NC ZF core. We propose that, in addition to the polycationic nature of the NC domain [72, 77, 79, 108], two motifs, one in the N-terminal 3_10_-helix and the other, an inverted motif in the NC-SP2 junction are responsible for NC-NA-NC and NA-NC-NA networks providing a source of quinary interactions. In the crowded in virio environment at neutral or mildly acidic pH, our model also involves quinary PR sequestration by the RNP, which dramatically enhances the global efficiency of the sequential cleavage. These findings are consistent with recent observations that HIV-1, and more broadly, retroviral NC phase separates in the intracellular environment [55].

**Fig 6.**
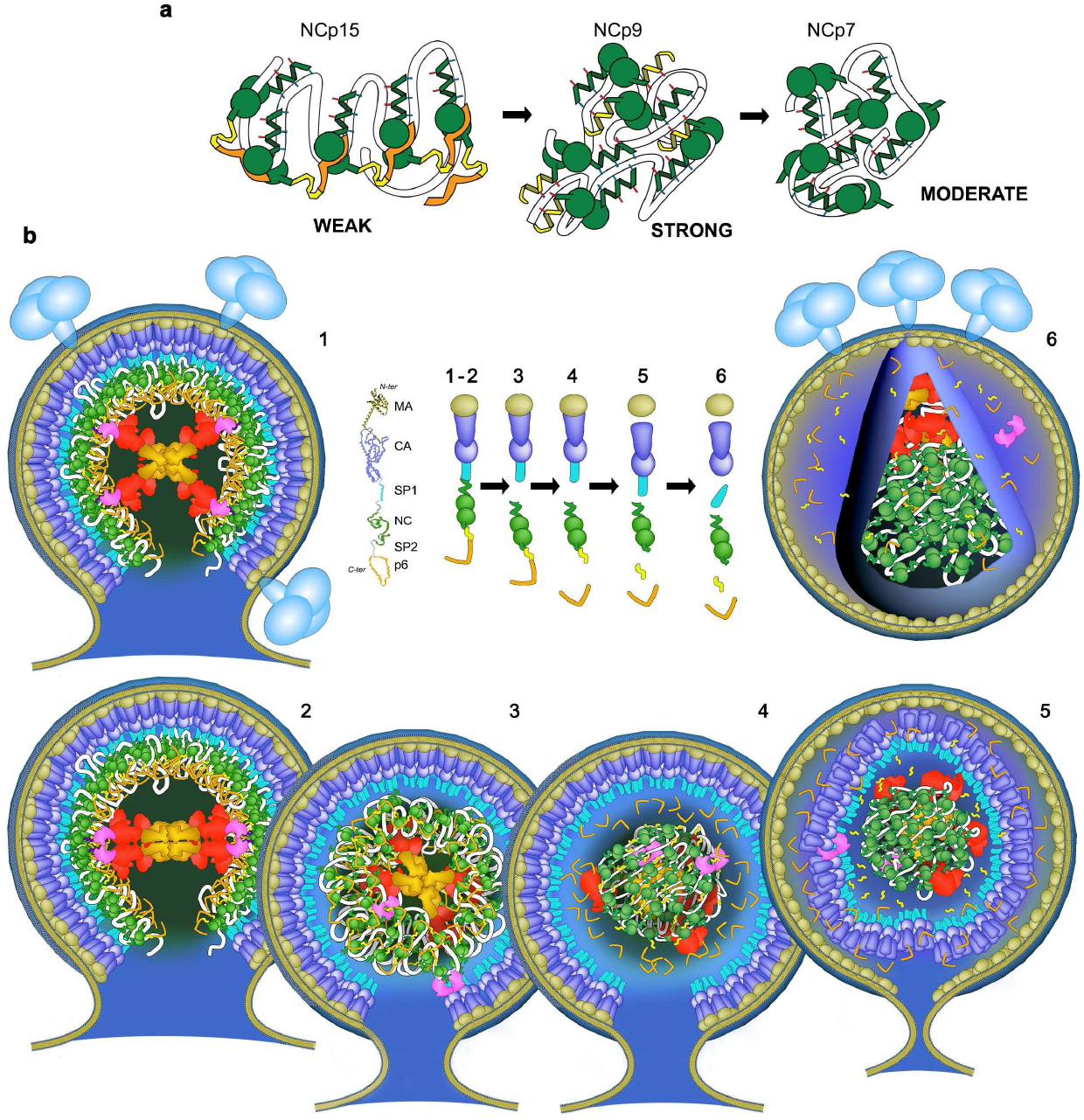
Nucleocapsid condensation, quinary interactions and WSM transition of the nucleocapsid during HIV maturation. a, Weak-Strong-Moderate quinary properties of NC proteins with RNA (white) throughout virus maturation. p6 (orange)-NCp7 (green) contacts assemble NCp15 networks upon the RNA, blocking exposed residues engaged in NCp9 or NCp7 : NA contacts. Cleavage of p6 unmasks the NCp9 domain, the NCp7-SP2 (yellow) domain engages NC:NA:NC and NA:NC:NA networks favouring strong RNA-mediated quinary interactions and RNP granulation. Cleavage of SP2 removes one “K/F/Q” NA binding patch (see Fig 2a), which reduces the quinary network between RNA and NCp7. b, Nucleocapsid WSM transition in the context of viral particle. (1) Virus particle at the plasma membrane bud from self-assembly of Gag and GagPol on gRNA, with the NC domain as the RNA binder. The gRNA is shown in white within the layer of NC domains, assembled as bundles of six at the base of Gag hexamers. (2) The dimerization of GagPol self-activates PR (pink) and initiates maturation. (3) The SP1-NC site is cleaved, liberating the gRNA:NCp15 RNP, which in turns sequesters PR. (4) Rapid cleavage of NCp15 into NCp9, which liberates p6, unlocks the strong quinary properties of NCp9. This quickly compacts the gRNA and favours viral budding. gRNA condensation allows internal reorganization of RT (red) and IN (gold), self-assembly of the conical capsid after separation of MA from CA-SP1 and maturation of CA-SP1 while NCp9 are matured to NCp7 in (5 and 6).

Our data confirm, first, that RNA-bound NCp15 avoids active RNP condensation within the NCp15-gRNA intermediate assembly. The intrinsically disordered p6 likely directs a quinary RNA-NCp15 network via NC:p6 intermolecular contacts and weakens the quinary RNA-NC interactions. Such assembly is deficient in actively aggregating within the viral core, while it should allow the 60 PR available in the particle to efficiently access the 2400 SP2-p6 cleavage sites and to jump from site to site within close distances (Figure 6). PR previously was shown to be activated by interacting with RNA [64]. We demonstrate here that fast PR turnover requires RNA-clustered NCp15 for the enzyme to be sequestrated, enhancing its local concentration and allowing it to efficiently cleave the SP2-p6 octapeptide sites.

We suggest the mechanistic origin of this sequestration to be the dynamic formation of a network of quinary interactions, derived from multiple, weakly attractive ternary contacts of PR-(SP2-p6)-RNA. Whilst the SP2-p6 octapeptide does not directly appear in contact with RNA, the exposed PR basic residues proposed to bind to RNA [64] should allow this ternary contact, whereas RNA-bound NCp15 should expose its SP2-p6 cleavage site to RNA-bound PR with the NC-SP2 site being inaccessible. One quinary effect would be that progressive cleavage of the SP2-p6 sites within the RNP results in loss of these ternary contacts, driving the change in enzyme absorption equilibrium – thus expelling PR from the RNP. Following cleavage of an SP2-p6 site, another effect may be a directed, rather than diffusive, propagation of PR. Inaccessibility of NC-SP2 implies a discrimination process that favors trans PR transfer to neighboring NCp15 molecules rather than cis sliding along the NCp9 moiety to cleave the RNA-bound NC-SP2 site, even though only 16 aa sequentially separate the two sites. Cleaved p6, already in contact with a neighboring NC domain, likely directs the trans event. Both effects are complementary to each other and consistent with our model.

When NCp9 forms, the NC-SP2 octapeptide site should bind more easily to RNA due to p6 separation, promoting both the PR recruitment for the NC-SP2 cleavage and the RNA condensation in concert with the other critical NC residues. Again a ternary PR-NCp9-RNA complex would be required, as the free NCp9 appeared here as a poor substrate for PR, confirming previous results [68]. We propose that the NC-SP2 site is in direct contact with RNA by way of the QxxFxxK triad. The critical NC-SP2 octapeptide also binds within the PR internal catalytic site for cleavage: this cleavage should require displacement of the bound RNA upon the surface of PR allowing a quinary contact between RNA and PR. In addition, longer RNA segments are required to strongly enhance NCp9 cleavage suggesting that PR transfers directly from cleaved NCp9 (i.e. NCp7) to neighboring NCp9. RNA condensation does not seem to provide any critical physical barrier to PR as NCp9 processing appears quite similar whether the reaction starts from NCp15 or NCp9 (Figure 3a). This suggests a perfect fit for ordered turnover by PR from site to site with two distinct waves, NCp15-to-NCp9 followed by NCp9-to-NCp7.

Whilst RNA and NC are dispensable for HIV-1 capsid assembly [115], integrative biochemical reconstitution studies have shown that RNA and other cofactors play important synergestic roles in HIV-1 assembly kinetics [116]. In our HIV-1 model, the granulation dynamics shown here opens new avenues at the mesoscopic scale to better understand how the surrounding capsid progresses towards a conical reassembly and what the implication and relocation of RT and IN proteins are. While both mature during GagPol processing, IN has been shown recently to be a key actor in properly coordinating RNP granular condensation and capsid reassembly within the viral particle [117–119]. These dynamics also offer new schemes to revisit the proposed implications of HIV-1 Nef, Vif, Tat and Vpr auxiliary proteins within the design of an infectious particle [120]. Our findings suggest tight temporal control of GagPol incorporation and PR auto-processing supported by the NC domain in GagPol during particle formation [121, 122]: this domain should help to position GagPol in the Gag assembly, facilitating the processed PR directed by the gRNA to digest the Gag-domains NC, SP2, p6. As highlighted by an independent study looking for production of GagPol VLPs without any viral accessory genes [123], the concomitant interactions of the exposed p6 domains with the ESCRT components, ALIX and more especially TSG101 [82, 124, 125], tightly coordinate complete processing of Gag within just a few minutes of particle release. Such proteins may also interact with the p6 domain of Gag - leading to weakening of the p6-RNA interaction, consistent with our model and in accordance with what we show here, namely, the exclusive release and processing of mature HIV-1 particles, including a condensed nucleocapsid, accelerated to the same time frame due to a gel phase separated effect.

Quinary interactions undoubtedly offer a missing link between molecular and cellular biology [17, 25, 28, 29] both in terms of fundamental understanding and therapeutic application. The quinary aspect of NC-RNA interactions regulating a biomolecular condensate, shown here, provide new perspectives for pharmacological targeting during particle production [126]. Finally, our HIV-1 model adds a novel dimension to the study of biomolecular condensates (BCs) and liquid-liquid phase separation (LLPS). Not only does it suggest concentration-driven accelerated enzyme activity in such BCs but that the cooperation of RNA, RNA-binding proteins and an embedded proteolytic machinery can create a scaffolding role for crowded architectures that dynamically regulate their own condensation through adjustment of quinary interactions [25, 29, 127]. Given that RNA-containing membraneless compartments have been linked to origins of life chemistry [128], such dynamic condensate regulation mechanisms may be universal properties of RNPs and, moreover, may have emerged early in the evolution of life.

## Supplementary Information

**Supplementary Figure 1** Large-scale NA binding of NC proteins followed by EMSA

**Supplementary Figure 2** Binding, condensation or aggregation of M13 ssDNA by NC followed by AFM

**Supplementary Figure 3** The PR-driven cleavage of NCp in vitro is strongly activated by NA

**Supplementary Figure 4** PR activation of NC proteolysis is modulated by NA length and NC:NAinteractions

**Supplementary Figure 5** PR is sequestered by the NCp15:ssDNA NP complex

**Supplementary Figure 6** Nucleocapsid condensation is concomitant with budding

**Supplementary Table 1** Kinetic parameters for RNP-modulated two-substrate model

**Supplementary Note** A polymer model of nucleic acid-modulated enzyme kinetics

1: Theoretical development

2: Computational implementation and model parameters

3: Analysis of model features

**Supplementary Appendix**

## Author Contributions

S. L. and G. M. conceived and designed the experiments. S. L. and T. E. performed the DNA mobility shift assays. S. L. did the AFM experiments. J. O. and S. L. did the DLS experiments. L. D., T. E., X. T., M. O.-O., M. R.-R. did the protease experiments. S.K. S. conceived and developed the theory, designed and implemented the kinetic model and the molecular dynamics simulations. R. J. G. and C. T. prepared NC proteins. L. D. and M. R.-R. prepared PR proteins. J.-C. P. and R. M. provided the collection of HIV-1 RNA fragments. R. J. G., C. L.-O. and N. G. did the virological and TEM experiments, with J. M. G. and A. M. advising. S. L., S.K. S. and G. M. analyzed the data and wrote the manuscript with strong support of R. J. G. and input from J. M. G., A. M., J.-C. P., R. M. and C. T. Figures were developed and arranged by S. L., G. M. and S. K. S. S.L. realized the illustrations.

## Acknowledegments

We thank Carlo Carolis, Biomolecular Screening and Protein Technology Unit (CRG), Carmen Lopez Iglesias and the TEM-SEM facilities of Science and Technical Centers of the Universitat de Barcelona (CCiT-UB), Cathy V. Hixson and Donald G. Johnson of Leidos Biomedical Research, Inc. and Eric Le Cam (IGR and CNRS). his work was supported in part by the European Project THINPAD “Targeting the HIV-1 Nucleocapsid Protein to fight Antiretroviral Drug Resistance” (FP7-Grant Agreement 601969), by Foundation Clinic, by ANRS, by SIDACTION, and with Federal funds from the NCI/NIH, under Contract No. HHSN261200800001E with Leidos Biomedical Research, Inc. (R.J.G.). S.L. acknowledges funding by the Marie-Curie IEF fellowship (FP7-Grant Agreement 237738) and is grateful to Maria Sol`a (IBMB-CSIC). S.K.S. and A.M. acknowledge support from amfAR Mathilde Krim Fellowship in Basic Biomedical Research number 108680 and the Spanish Ministry of Economy and Competitiveness and FEDER (Grant no. SAF2013-46077-R). S.K.S. also gratefully acknowledges support from the Volkswagen Foundation “Experiment! Funding Initiative” grant number 93874 and from the Klaus Tschira Stiftung.

## References

1. André AA, Spruijt E. Liquid–liquid phase separation in crowded environments. International Journal of Molecular Sciences. 2020;21(16):5908.

2. Yoshizawa T, Nozawa RS, Jia TZ, Saio T, Mori E. Biological phase separation: cell biology meets biophysics. Biophysical reviews. 2020;12(2):519–539.

3. Alberti S, Hyman AA. Biomolecular condensates at the nexus of cellular stress, protein aggregation disease and ageing. Nature Reviews Molecular Cell Biology. 2021;22(3):196–213.

4. Alberti S, Gladfelter A, Mittag T. Considerations and challenges in studying liquid-liquid phase separation and biomolecular condensates. Cell. 2019;176(3):419–434.

5. Feng Z, Chen X, Wu X, Zhang M. Formation of biological condensates via phase separation: Characteristics, analytical methods, and physiological implications. Journal of Biological Chemistry. 2019;294(40):14823–14835.

6. Wallace EW, Kear-Scott JL, Pilipenko EV, Schwartz MH, Laskowski PR, Rojek AE, et al. Reversible, specific, active aggregates of endogenous proteins assemble upon heat stress. Cell. 2015;162(6):1286–1298.

7. Franzmann TM, Alberti S. Protein phase separation as a stress survival strategy. Cold Spring Harbor perspectives in biology. 2019;11(6):a034058.

8. Panas MD, Ivanov P, Anderson P. Mechanistic insights into mammalian stress granule dynamics. Journal of Cell Biology. 2016;215(3):313–323.

9. Riback JA, Katanski CD, Kear-Scott JL, Pilipenko EV, Rojek AE, Sosnick TR, et al. Stress-triggered phase separation is an adaptive, evolutionarily tuned response. Cell. 2017;168(6):1028–1040.

10. Ambadipudi S, Biernat J, Riedel D, Mandelkow E, Zweckstetter M. Liquid–liquid phase separation of the microtubule-binding repeats of the Alzheimer-related protein Tau. Nature communications. 2017;8(1):1–13.

11. Cai D, Liu Z, Lippincott-Schwartz J. Biomolecular Condensates and Their Links to Cancer Progression. Trends in biochemical sciences. 2021;.

12. Nakashima KK, Vibhute MA, Spruijt E. Biomolecular chemistry in liquid phase separated compartments. Frontiers in molecular biosciences. 2019;6:21.

13. O’Flynn BG, Mittag T. The role of liquid–liquid phase separation in regulating enzyme activity. Current opinion in cell biology. 2021;69:70–79.

14. Banani SF, Lee HO, Hyman AA, Rosen MK. Biomolecular condensates: organizers of cellular biochemistry. Nature reviews Molecular cell biology. 2017;18(5):285–298.

15. Laflamme G, Mekhail K. Biomolecular condensates as arbiters of biochemical reactions inside the nucleus. Communications Biology. 2020;3(1):1–8.

16. Zhang Y, Narlikar GJ, Kutateladze TG. Enzymatic reactions inside biological condensates. Journal of molecular biology. 2021;433(12):166624.

17. Buchan JR. mRNP granules: assembly, function, and connections with disease. RNA biology. 2014;11(8):1019–1030.

18. Roden C, Gladfelter AS. RNA contributions to the form and function of biomolecular condensates. Nature Reviews Molecular Cell Biology. 2021;22(3):183–195.

19. Sanders DW, Kedersha N, Lee DS, Strom AR, Drake V, Riback JA, et al. Competing protein-RNA interaction networks control multiphase intracellular organization. Cell. 2020;181(2):306–324.

20. Sanulli S, Narlikar GJ. Generation and Biochemical Characterization of Phase-Separated Droplets Formed by Nucleic Acid Binding Proteins: Using HP1 as a Model System. Current Protocols. 2021;1(5):e109.

21. Luo J, Qu L, Gao F, Lin J, Liu J, Lin A. LncRNAs: Architectural Scaffolds or More Potential Roles in Phase Separation. Frontiers in Genetics. 2021;12:369.

22. Louka A, Zacco E, Temussi PA, Tartaglia GG, Pastore A. RNA as the stone guest of protein aggregation. Nucleic Acids Research. 2020;48(21):11880–11889.

23. Wiedner HJ, Giudice J. It’s not just a phase: function and characteristics of RNA-binding proteins in phase separation. Nature Structural & Molecular Biology. 2021;28(6):465–473.

24. Gotor NL, Armaos A, Calloni G, Torrent Burgas M, Vabulas RM, De Groot NS, et al. RNA-binding and prion domains: the Yin and Yang of phase separation. Nucleic acids research. 2020;48(17):9491–9504.

25. Guo L, Shorter J. It’s raining liquids: RNA tunes viscoelasticity and dynamics of membraneless organelles. Molecular cell. 2015;60(2):189–192.

26. Woodruff JB, Hyman AA, Boke E. Organization and function of non-dynamic biomolecular condensates. Trends in biochemical sciences. 2018;43(2):81–94.

27. Patel A, Lee HO, Jawerth L, Maharana S, Jahnel M, Hein MY, et al. A liquid-to-solid phase transition of the ALS protein FUS accelerated by disease mutation. Cell. 2015;162(5):1066–1077.

28. Chien P, Gierasch LM. Challenges and dreams: physics of weak interactions essential to life. Molecular biology of the cell. 2014;25(22):3474–3477.

29. McConkey EH. Molecular evolution, intracellular organization, and the quinary structure of proteins. Proceedings of the National Academy of Sciences. 1982;79(10):3236–3240.

30. Monteith WB, Cohen RD, Smith AE, Guzman-Cisneros E, Pielak GJ. Quinary structure modulates protein stability in cells. Proceedings of the National Academy of Sciences. 2015;112(6):1739–1742.

31. Guin D, Gruebele M. Weak chemical interactions that drive protein evolution: crowding, sticking, and quinary structure in folding and function. Chemical reviews. 2019;119(18):10691–10717.

32. Rickard MM, Zhang Y, Gruebele M, Pogorelov TV. In-Cell Protein–Protein Contacts: Transient Interactions in the Crowd. The journal of physical chemistry letters. 2019;10(18):5667–5673.

33. Gopi S, Naganathan AN. Non-specific DNA-driven quinary interactions promote structural transitions in proteins. Physical Chemistry Chemical Physics. 2020;22(22):12671–12677.

34. Ziegler SJ, Mallinson SJ, John PCS, Bomble YJ. Advances in integrative structural biology: Towards understanding protein complexes in their cellular context. Computational and Structural Biotechnology Journal. 2020;.

35. Li P, Banjade S, Cheng HC, Kim S, Chen B, Guo L, et al. Phase transitions in the assembly of multivalent signalling proteins. Nature. 2012;483(7389):336–340.

36. Choi JM, Holehouse AS, Pappu RV. Physical principles underlying the complex biology of intracellular phase transitions. Annual Review of Biophysics. 2020;49:107–133.

37. Dignon GL, Best RB, Mittal J. Biomolecular phase separation: From molecular driving forces to macroscopic properties. Annual review of physical chemistry. 2020;71:53–75.

38. Ghosh A, Mazarakos K, Zhou HX. Three archetypical classes of macromolecular regulators of protein liquid–liquid phase separation. Proceedings of the National Academy of Sciences. 2019;116(39):19474–19483.

39. Nott TJ, Petsalaki E, Farber P, Jervis D, Fussner E, Plochowietz A, et al. Phase transition of a disordered nuage protein generates environmentally responsive membraneless organelles. Molecular cell. 2015;57(5):936–947.

40. Zhou HX, Nguemaha V, Mazarakos K, Qin S. Why do disordered and structured proteins behave differently in phase separation? Trends in biochemical sciences. 2018;43(7):499–516.

41. Bratek-Skicki A, Pancsa R, Meszaros B, Van Lindt J, Tompa P. A guide to regulation of the formation of biomolecular condensates. The FEBS journal. 2020;287(10):1924–1935.

42. Van Lindt J, Bratek-Skicki A, Nguyen PN, Pakravan D, Durán-Armenta LF, Tantos A, et al. A generic approach to study the kinetics of liquid–liquid phase separation under near-native conditions. Communications Biology. 2021;4(1):1–8.

43. Sanchez-Burgos I, Espinosa JR, Joseph JA, Collepardo-Guevara R. Valency and binding affinity variations can regulate the multilayered organization of protein condensates with many components. Biomolecules. 2021;11(2):278.

44. Espinosa JR, Joseph JA, Sanchez-Burgos I, Garaizar A, Frenkel D, Collepardo-Guevara R. Liquid network connectivity regulates the stability and composition of biomolecular condensates with many components. Proceedings of the National Academy of Sciences. 2020;117(24):13238–13247.

45. Dar F, Pappu R. Phase separation: Restricting the sizes of condensates. Elife. 2020;9:e59663.

46. Monahan Z, Ryan VH, Janke AM, Burke KA, Rhoads SN, Zerze GH, et al. Phosphorylation of the FUS low-complexity domain disrupts phase separation, aggregation, and toxicity. The EMBO journal. 2017;36(20):2951–2967.

47. Hofweber M, Hutten S, Bourgeois B, Spreitzer E, Niedner-Boblenz A, Schifferer M, et al. Phase separation of FUS is suppressed by its nuclear import receptor and arginine methylation. Cell. 2018;173(3):706–719.

48. Qamar S, Wang G, Randle SJ, Ruggeri FS, Varela JA, Lin JQ, et al. FUS phase separation is modulated by a molecular chaperone and methylation of arginine cation-π interactions. Cell. 2018;173(3):720–734.

49. Etibor TA, Yamauchi Y, Amorim MJ. Liquid Biomolecular Condensates and Viral Lifecycles: Review and Perspectives. Viruses. 2021;13(3):366.

50. Brocca S, Grandori R, Longhi S, Uversky V. Liquid–liquid phase separation by intrinsically disordered protein regions of viruses: Roles in viral life cycle and control of virus–host interactions. International Journal of Molecular Sciences. 2020;21(23):9045.

51. Iserman C, Roden CA, Boerneke MA, Sealfon RS, McLaughlin GA, Jungreis I, et al. Genomic RNA elements drive phase separation of the SARS-CoV-2 nucleocapsid. Molecular cell. 2020;80(6):1078–1091.

52. Chen H, Cui Y, Han X, Hu W, Sun M, Zhang Y, et al. Liquid–liquid phase separation by SARS-CoV-2 nucleocapsid protein and RNA. Cell research. 2020;30(12):1143–1145.

53. Savastano A, de Opakua AI, Rankovic M, Zweckstetter M. Nucleocapsid protein of SARS-CoV-2 phase separates into RNA-rich polymerase-containing condensates. Nature communications. 2020;11(1):1–10.

54. Perdikari TM, Murthy AC, Ryan VH, Watters S, Naik MT, Fawzi NL. SARS-CoV-2 nucleocapsid protein phase-separates with RNA and with human hnRNPs. The EMBO journal. 2020;39(24):e106478.

55. Monette A, Niu M, Chen L, Rao S, Gorelick RJ, Mouland AJ. Pan-retroviral nucleocapsid-mediated phase separation regulates genomic RNA positioning and trafficking. Cell reports. 2020;31(3):107520.

56. Monette A, Mouland AJ. Zinc and copper ions differentially regulate prion-like phase separation dynamics of pan-virus nucleocapsid biomolecular condensates. Viruses. 2020;12(10):1179.

57. Mirambeau G, Lyonnais S, Coulaud D, Hameau L, Lafosse S, Jeusset J, et al. Transmission electron microscopy reveals an optimal HIV-1 nucleocapsid aggregation with single-stranded nucleic acids and the mature HIV-1 nucleocapsid protein. Journal of molecular biology. 2006;364(3):496–511.

58. Mirambeau G, Lyonnais S, Coulaud D, Hameau L, Lafosse S, Jeusset J, et al. HIV-1 protease and reverse transcriptase control the architecture of their nucleocapsid partner. PloS one. 2007;2(8):e669.

59. Mirambeau G, Lyonnais S, Gorelick RJ. Features, processing states, and heterologous protein interactions in the modulation of the retroviral nucleocapsid protein function. RNA biology. 2010;7(6):724–734.

60. Wang W, Naiyer N, Mitra M, Li J, Williams MC, Rouzina I, et al. Distinct nucleic acid interaction properties of HIV-1 nucleocapsid protein precursor NCp15 explain reduced viral infectivity. Nucleic acids research. 2014;42(11):7145–7159.

61. Sundquist WI, Kräusslich HG. HIV-1 assembly, budding, and maturation. Cold Spring Harbor perspectives in medicine. 2012;2(7):a006924.

62. Konvalinka J, Kräusslich HG, Müller B. Retroviral proteases and their roles in virion maturation. Virology. 2015;479:403–417.

63. Deshmukh L, Ghirlando R, Clore GM. Conformation and dynamics of the Gag polyprotein of the human immunodeficiency virus 1 studied by NMR spectroscopy. Proceedings of the National Academy of Sciences. 2015;112(11):3374–3379.

64. Potempa M, Nalivaika E, Ragland D, Lee SK, Schiffer CA, Swanstrom R. A direct interaction with RNA dramatically enhances the catalytic activity of the HIV-1 protease in vitro. Journal of molecular biology. 2015;427(14):2360–2378.

65. de Marco A, Heuser AM, Glass B, Kräusslich HG, Müller B, Briggs JA. Role of the SP2 domain and its proteolytic cleavage in HIV-1 structural maturation and infectivity. Journal of virology. 2012;86(24):13708–13716.

66. Wu T, Gorelick RJ, Levin JG. Selection of fully processed HIV-1 nucleocapsid protein is required for optimal nucleic acid chaperone activity in reverse transcription. Virus research. 2014;193:52–64.

67. Todd MJ, Semo N, Freire E. The structural stability of the HIV-1 protease. Journal of molecular biology. 1998;283(2):475–488.

68. Pettit SC, Sheng N, Tritch R, Erickson-Viitanen S, Swanstrom R. The regulation of sequential processing of HIV-1 Gag by the viral protease. In: Aspartic Proteinases. Springer; 1998. p. 15–25.

69. Sheng N, Pettit SC, Tritch RJ, Ozturk DH, Rayner MM, Swanstrom R, et al. Determinants of the human immunodeficiency virus type 1 p15NC-RNA interaction that affect enhanced cleavage by the viral protease. Journal of virology. 1997;71(8):5723–5732.

70. Amarasinghe GK, De Guzman RN, Turner RB, Chancellor KJ, Wu ZR, Summers MF. NMR structure of the HIV-1 nucleocapsid protein bound to stem-loop SL2 of the Ψ-RNA packaging signal. Implications for genome recognition. Journal of molecular biology. 2000;301(2):491–511.

71. De Guzman RN, Wu ZR, Stalling CC, Pappalardo L, Borer PN, Summers MF. Structure of the HIV-1 nucleocapsid protein bound to the SL3 Ψ-RNA recognition element. Science. 1998;279(5349):384–388.

72. Wu H, Mitra M, Naufer MN, McCauley MJ, Gorelick RJ, Rouzina I, et al. Differential contribution of basic residues to HIV-1 nucleocapsid protein’s nucleic acid chaperone function and retroviral replication. Nucleic acids research. 2014;42(4):2525–2537.

73. Le Cam E, Coulaud D, Delain E, Petitjean P, Roques BP, Gérard D, et al. Properties and growth mechanism of the ordered aggregation of a model RNA by the HIV-1 nucleocapsid protein: An electron microscopy investigation. Biopolymers: Original Research on Biomolecules. 1998;45(3):217–229.

74. Mouhand A, Pasi M, Catala M, Zargarian L, Belfetmi A, Barraud P, et al. Overview of the nucleic-acid binding properties of the HIV-1 nucleocapsid protein in its different maturation states. Viruses. 2020;12(10):1109.

75. Retureau R, Oguey C, Mauffret O, Hartmann B. Structural Explorations of NCp7–Nucleic Acid Complexes Give Keys to Decipher the Binding Process. Journal of molecular biology. 2019;431(10):1966–1980.

76. Mouhand A, Belfetmi A, Catala M, Larue V, Zargarian L, Brachet F, et al. Modulation of the HIV nucleocapsid dynamics finely tunes its RNA-binding properties during virion genesis. Nucleic acids research. 2018;46(18):9699–9710.

77. Khan R, Giedroc DP. Nucleic acid binding properties of recombinant Zn2 HIV-1 nucleocapsid protein are modulated by COOH-terminal processing. Journal of Biological Chemistry. 1994;269(36):22538–22546.

78. Wu H, Rouzina I, Williams MC. Single-molecule stretching studies of RNA chaperones. RNA biology. 2010;7(6):712–723.

79. Cruceanu M, Urbaneja MA, Hixson CV, Johnson DG, Datta SA, Fivash MJ, et al. Nucleic acid binding and chaperone properties of HIV-1 Gag and nucleocapsid proteins. Nucleic acids research. 2006;34(2):593–605.

80. Lee SK, Potempa M, Swanstrom R. The choreography of HIV-1 proteolytic processing and virion assembly. Journal of Biological Chemistry. 2012;287(49):40867–40874.

81. Könnyű B, Sadiq SK, Turányi T, Hírmondó R, Müller B, Kräusslich HG, et al. Gag-Pol processing during HIV-1 virion maturation: a systems biology approach. PLoS computational biology. 2013;9(6):e1003103.

82. Dussupt V, Sette P, Bello NF, Javid MP, Nagashima K, Bouamr F. Basic residues in the nucleocapsid domain of Gag are critical for late events of HIV-1 budding. Journal of virology. 2011;85(5):2304–2315.

83. Moore MD, Fu W, Soheilian F, Nagashima K, Ptak RG, Pathak VK, et al. Sub-optimal inhibition of protease activity in human immunodeficiency virus type 1: effects on virion morphogenesis and RNA maturation. Virology. 2008;379(1):152–160.

84. Coren LV, Thomas JA, Chertova E, Sowder RC, Gagliardi TD, Gorelick RJ, et al. Mutational analysis of the C-terminal gag cleavage sites in human immunodeficiency virus type 1. Journal of virology. 2007;81(18):10047–10054.

85. Guo J, Wu T, Anderson J, Kane BF, Johnson DG, Gorelick RJ, et al. Zinc finger structures in the human immunodeficiency virus type 1 nucleocapsid protein facilitate efficient minus-and plus-strand transfer. Journal of virology. 2000;74(19):8980–8988.

86. Stewart-Maynard KM, Cruceanu M, Wang F, Vo MN, Gorelick RJ, Williams MC, et al. Retroviral nucleocapsid proteins display nonequivalent levels of nucleic acid chaperone activity. Journal of virology. 2008;82(20):10129–10142.

87. Carteau S, Gorelick RJ, Bushman FD. Coupled integration of human immunodeficiency virus type 1 cDNA ends by purified integrase in vitro: stimulation by the viral nucleocapsid protein. Journal of virology. 1999;73(8):6670–6679.

88. Wu H, Mitra M, McCauley MJ, Thomas JA, Rouzina I, Musier-Forsyth K, et al. Aromatic residue mutations reveal direct correlation between HIV-1 nucleocapsid protein’s nucleic acid chaperone activity and retroviral replication. Virus research. 2013;171(2):263–277.

89. Billich A, Hammerschmid F, Winkler G. Purification, assay and kinetic features of HIV-1 proteinase. Biol Chem Hoppe Seyler. 1990;371:265–272.

90. Bannwarth L, Rose T, Dufau L, Vanderesse R, Dumond J, Jamart-Grégoire B, et al. Dimer disruption and monomer sequestration by alkyl tripeptides are successful strategies for inhibiting wild-type and multidrug-resistant mutated HIV-1 proteases. Biochemistry. 2009;48(2):379–387.

91. Davis DA, Brown CA, Newcomb FM, Boja ES, Fales HM, Kaufman J, et al. Reversible oxidative modification as a mechanism for regulating retroviral protease dimerization and activation. Journal of virology. 2003;77(5):3319–3325.

92. Rothemund PW. Folding DNA to create nanoscale shapes and patterns. Nature. 2006;440(7082):297–302.

93. Sinck L, Richer D, Howard J, Alexander M, Purcell DF, Marquet R, et al. In vitro dimerization of human immunodeficiency virus type 1 (HIV-1) spliced RNAs. Rna. 2007;13(12):2141–2150.

94. Abd El-Wahab EW, Smyth RP, Mailler E, Bernacchi S, Vivet-Boudou V, Hijnen M, et al. Specific recognition of the HIV-1 genomic RNA by the Gag precursor. Nature communications. 2014;5(1):1–13.

95. Goldschmidt V, Paillart JC, Rigourd M, Ehresmann B, Aubertin AM, Ehresmann C, et al. Structural variability of the initiation complex of HIV-1 reverse transcription. Journal of Biological Chemistry. 2004;279(34):35923–35931.

96. Hamon L, Pastre D, Dupaigne P, Breton CL, Cam EL, Pietrement O. High-resolution AFM imaging of single-stranded DNA-binding (SSB) protein—DNA complexes. Nucleic acids research. 2007;35(8):e58.

97. Gonda MA, Aaronson SA, Ellmore N, Zeve VH, Nagashima K. Ultrastructural studies of surface features of human normal and tumor cells in tissue culture by scanning and transmission electron microscopy. Journal of the National Cancer Institute. 1976;56(2):245–263.

98. Ott DE, Chertova EN, Busch LK, Coren LV, Gagliardi TD, Johnson DG. Mutational analysis of the hydrophobic tail of the human immunodeficiency virus type 1 p6Gag protein produces a mutant that fails to package its envelope protein. Journal of virology. 1999;73(1):19–28.

99. Clouse KA, Powell D, Washington I, Poli G, Strebel K, Farrar W, et al. Monokine regulation of human immunodeficiency virus-1 expression in a chronically infected human T cell clone. The Journal of Immunology. 1989;142(2):431–438.

100. Lopez-Iglesias C, Puvion-Dutilleul F. Visualization of glycoproteins after tunicamycin and monensin treatment of herpes simplex virus infected cells. Journal of ultrastructure and molecular structure research. 1988;101(1):75–91.

101. Sadiq SK, Coveney PV. Computing the role of near attack conformations in an enzyme-catalyzed nucleophilic bimolecular reaction. Journal of chemical theory and computation. 2015;11(1):316–324.

102. Prabu-Jeyabalan M, Nalivaika EA, King NM, Schiffer CA. Structural basis for coevolution of a human immunodeficiency virus type 1 nucleocapsid-p1 cleavage site with a V82A drug-resistant mutation in viral protease. Journal of virology. 2004;78(22):12446–12454.

103. Duan Y, Wu C, Chowdhury S, Lee MC, Xiong G, Zhang W, et al. A point-charge force field for molecular mechanics simulations of proteins based on condensed-phase quantum mechanical calculations. Journal of computational chemistry. 2003;24(16):1999–2012.

104. Harvey MJ, Giupponi G, Fabritiis GD. ACEMD: accelerating biomolecular dynamics in the microsecond time scale. Journal of chemical theory and computation. 2009;5(6):1632–1639.

105. Vo MN, Barany G, Rouzina I, Musier-Forsyth K. Effect of Mg2+ and Na+ on the nucleic acid chaperone activity of HIV-1 nucleocapsid protein: implications for reverse transcription. Journal of molecular biology. 2009;386(3):773–788.

106. Williams MC, Rouzina I, Wenner JR, Gorelick RJ, Musier-Forsyth K, Bloomfield VA. Mechanism for nucleic acid chaperone activity of HIV-1 nucleocapsid protein revealed by single molecule stretching. Proceedings of the National Academy of Sciences. 2001;98(11):6121–6126.

107. Fisher RJ, Fivash MJ, Stephen AG, Hagan NA, Shenoy SR, Medaglia MV, et al. Complex interactions of HIV-1 nucleocapsid protein with oligonucleotides. Nucleic acids research. 2006;34(2):472–484.

108. Qualley DF, Stewart-Maynard KM, Wang F, Mitra M, Gorelick RJ, Rouzina I, et al. C-terminal domain modulates the nucleic acid chaperone activity of human T-cell leukemia virus type 1 nucleocapsid protein via an electrostatic mechanism. Journal of Biological Chemistry. 2010;285(1):295–307.

109. Sadiq SK, Könnyű B, Müller V, Coveney PV. Reaction kinetics of catalyzed competitive heteropolymer cleavage. The Journal of Physical Chemistry B. 2011;115(37):11017–11027.

110. Venken T, Voet A, De Maeyer M, De Fabritiis G, Sadiq SK. Rapid conformational fluctuations of disordered HIV-1 fusion peptide in solution. Journal of chemical theory and computation. 2013;9(7):2870–2874.

111. Roberts MM, Copeland TD, Oroszlan S. In situ processing of a retroviral nucleocapsid protein by the viral proteinase. Protein Engineering, Design and Selection. 1991;4(6):695–700.

112. Jouvenet N, Simon SM, Bieniasz PD. Visualizing HIV-1 assembly. Journal of molecular biology. 2011;410(4):501–511.

113. Votteler J, Sundquist WI. Virus budding and the ESCRT pathway. Cell host & microbe. 2013;14(3):232–241.

114. Prescher J, Baumgärtel V, Ivanchenko S, Torrano AA, Bräuchle C, Müller B, et al. Super-resolution imaging of ESCRT-proteins at HIV-1 assembly sites. PLoS pathogens. 2015;11(2):e1004677.

115. Mattei S, Flemming A, Anders-össwein M, Kräusslich HG, Briggs JA, Müller B. RNA and nucleocapsid are dispensable for mature HIV-1 capsid assembly. Journal of virology. 2015;89(19):9739–9747.

116. Kucharska I, Ding P, Zadrozny KK, Dick RA, Summers MF, Ganser-Pornillos BK, et al. Biochemical reconstitution of HIV-1 assembly and maturation. Journal of virology. 2020;94(5):e01844–19.

117. Fontana J, Jurado KA, Cheng N, Ly NL, Fuchs JR, Gorelick RJ, et al. Distribution and redistribution of HIV-1 nucleocapsid protein in immature, mature, and integrase-inhibited virions: a role for integrase in maturation. Journal of virology. 2015;89(19):9765–9780.

118. Kessl JJ, Kutluay SB, Townsend D, Rebensburg S, Slaughter A, Larue RC, et al. HIV-1 integrase binds the viral RNA genome and is essential during virion morphogenesis. Cell. 2016;166(5):1257–1268.

119. Elliott JL, Eschbach JE, Koneru PC, Li W, Puray-Chavez M, Townsend D, et al. Integrase-RNA interactions underscore the critical role of integrase in HIV-1 virion morphogenesis. Elife. 2020;9:e54311.

120. Cuccurullo EC, Valentini C, Pizzato M. Retroviral factors promoting infectivity. Progress in molecular biology and translational science. 2015;129:213–251.

121. Cen S, Niu M, Saadatmand J, Guo F, Huang Y, Nabel GJ, et al. Incorporation of pol into human immunodeficiency virus type 1 Gag virus-like particles occurs independently of the upstream Gag domain in Gag-pol. Journal of virology. 2004;78(2):1042–1049.

122. Figueiredo A, Moore KL, Mak J, Sluis-Cremer N, de Bethune MP, Tachedjian G. Potent nonnucleoside reverse transcriptase inhibitors target HIV-1 Gag-Pol. PLoS pathogens. 2006;2(11):e119.

123. Bendjennat M, Saffarian S. The race against protease activation defines the role of ESCRTs in HIV budding. PLoS pathogens. 2016;12(6):e1005657.

124. Chamontin C, Rassam P, Ferrer M, Racine PJ, Neyret A, Lainé S, et al. HIV-1 nucleocapsid and ESCRT-component Tsg101 interplay prevents HIV from turning into a DNA-containing virus. Nucleic acids research. 2015;43(1):336–347.

125. Popova E, Popov S, Göttlinger HG. Human immunodeficiency virus type 1 nucleocapsid p1 confers ESCRT pathway dependence. Journal of virology. 2010;84(13):6590–6597.

126. Mori M, Kovalenko L, Lyonnais S, Antaki D, Torbett BE, Botta M, et al. Nucleocapsid protein: a desirable target for future therapies against HIV-1. The Future of HIV-1 Therapeutics. 2015; p. 53–92.

127. Dumont S, Prakash M. Emergent mechanics of biological structures. Molecular Biology of the Cell. 2014;25(22):3461–3465.

128. Poudyal RR, Pir Cakmak F, Keating CD, Bevilacqua PC. Physical principles and extant biology reveal roles for RNA-containing membraneless compartments in origins of life chemistry. Biochemistry. 2018;57(17):2509–2519.

