## Supplementary Information for "The HIV-1 ribonucleoprotein dynamically regulates its condensate behavior and drives acceleration of protease activity through membraneless granular phase separation"

Sébastien Lyonnais[1,2,♯,*], S. Kashif Sadiq[3,4,5,♯,*], Cristina Lorca-Oró[1], Laure Dufau[6], Sara Nieto-Marquez[1], Tuixent Escriba[1], Natalia Gabrielli[1], Xiao Tan[1,6], Mohamed Ouizougun-Oubari[1], Josephine Okoronkwo[1], Michèle Reboud-Ravaux[6], José Maria Gatell[1,7], Roland Marquet[8], Jean-Christophe Paillart[8], Andreas Meyerhans[3,9], Carine Tisné[10], Robert J. Gorelick[11] and Gilles Mirambeau[1, 12,♯,*]

[1] Infectious disease & AIDS Research Unit, IDIBAPS, Villaroel 170, Barcelona, Spain.

[2] CEMIPAI, UAR 3725 Université de Montpellier - CNRS, 1919 Route de Mende, 34000 Montpellier.

[3] Infection Biology Laboratory, DCEXS, Universitat Pompeu Fabra, Barcelona, Spain.

[4] Molecular and Cellular Modeling Group, Heidelberg Institute for Theoretical Studies (HITS), Schloss-Wolfsbrunnenweg 35, 69118 Heidelberg, Germany

[5] Genome Biology Unit, European Molecular Biology Laboratory, Meyerhofstrasse 1, 69117 Heidelberg, Germany

[6] UMR/CNRS 8256, IBPS, Sorbonne Universités, UPMC Univ Paris 06, Paris, France.

[7] University of Barcelona, Santapau 62, 08016 Barcelona, Spain

[8] Université de Strasbourg, CNRS, Architecture et Réactivité de l'ARN, UPR 9002, 2 Allée Conrad Roentgen, F-67000 Strasbourg, France

[9] Institució Catalana de Recerca i Estudis Avançats (ICREA), Barcelona, Spain.

[10] Laboratoire de Cristallographie et RMN Biologiques, CNRS, Paris Sorbonne Cité, 4 Avenue de l'Observatoire, Paris, 75006, France.

[11] AIDS and Cancer Virus Program, Frederick National Laboratory for Cancer Research, Frederick, MD 21701, USA

[12] Sorbonne Université, Laboratoire Arago, UMR BIOM, Avenue Pierre Fabre, 66650 Banyuls-sur-mer, France

[♯] These authors contributed equally.

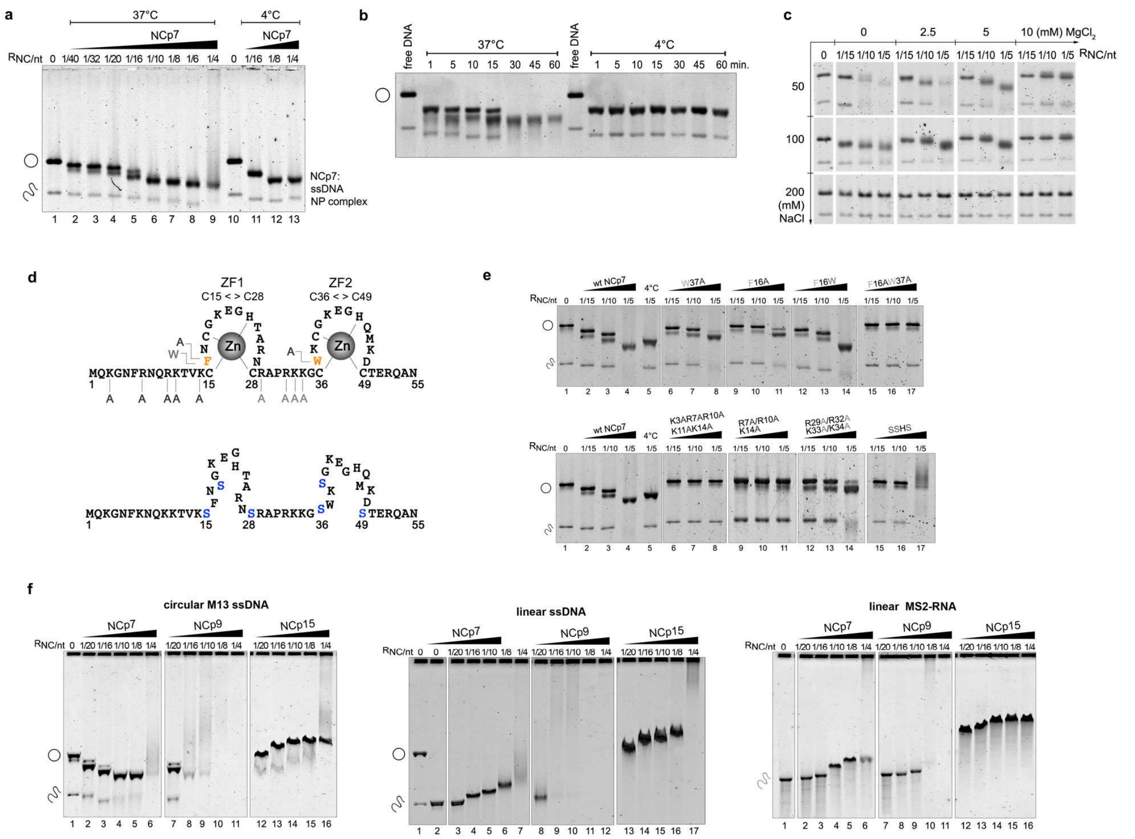


**Supplementary Figure 1 | Large-scale NA binding of NC proteins followed by EMSA. a,** The progressive condensation of M13 ssDNA by NCp7 is followed by stepwise addition of NCp7 at 37°C and 4°C. The NCp7/nt ratio are indicated and correspond, respectively (lanes 2 to 9) to 75, 93, 150, 166, 300, 375, 500 and 750 nM concentrations of NCp7. **b**, Kinetics of M13 ssDNA shows fast compaction and slow aggregation at 1 NCp7/8 nt ratio, at 37°C and 4°C in presence of 100 mM NaCl and 5 mM $MgCl_2$. **c**, Effect of monovalent ($Na^+$) and divalent (Mg2$^+$) cation addition on NCp7/M13 ssDNA interactions for three increasing NCp7 concentrations (200, 300 and 600 nM). The lowest monovalent salt concentration increased NCp7/ssDNA aggregation and a strong electrostatic competition was observed by both $Na^+$ or $Mg^+$, or in combination. **d,** Sequences of the NCp7 mutants. Amino acid replacements are indicated by a colour code reported in the EMSA experiments. The SSHS mutant substitutes serines for the cysteine residues that bind zinc to generate the apo-protein. **e**, M13 ssDNA was titrated using three increasing protein concentrations (200, 300 and 600 nM) in a buffer containing 100mM NaCl and 5mM $MgCl_2$. Binding reactions were carried out at 37°C for 30 min, except for the indicated lane at 4°C for a duplicate of the 1/5nt ratio with the wt NCp7. **f**, Titration of NCp7, NCp9 and NCp15 binding on M13 ssDNA (circular or linearized), or MS2-RNA (linear) in 100 mM NaCl and 2.5 mM $MgCl_2$ show differential compaction/binding/aggregation. Binding of NCp7 to linear M13 ssDNA or MS2 RNA produced NP complexes of lower mobility, which eventually aggregated, for lowest protein/nt ratios than with the circular ssDNA.

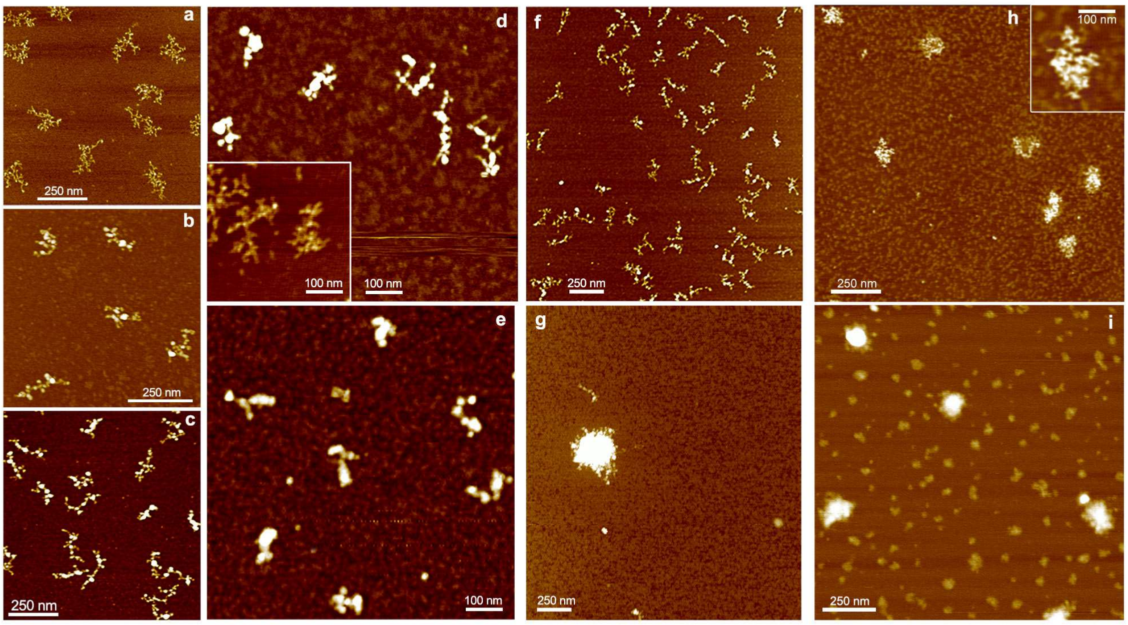


**Supplementary Figure 2 | Binding, condensation or aggregation of M13 ssDNA by NC followed by AFM. a,** free M13 ssDNA. **b, c, d,** Binding and progressive condensation of M13 ssDNA for 1 NCp7/20 nt; 1 NCp7/15 nt; 1 NCp7/10 nt, respectively. The insert in (**d**) is the free ssDNA at the same scale to appreciate the compaction and the melting of secondary structures by NCp7. **e,** NP complexes of maximum compaction for 1 NCp7/5 nt. **f,** field of individual and condensed NP complexes obtained at 1 NCp9/12 nt in 5 mM magnesium show NP complexes comparable to those obtained with NCp7. **g**, at R= 1 NCp9/8 nt., highly dense spheroids are formed, containing thousands of NCp9:ssDNA NP condensates joined together, while the mica surface appeared completely empty of individual complexes. **h, i,** The assembly of NCp15 on M13 ssDNA (1/20 nt. and 1/12 nt., respectively) show more passive NA binding without the DNA backbone bridging characteristic of NCp7 and NCp9 binding.

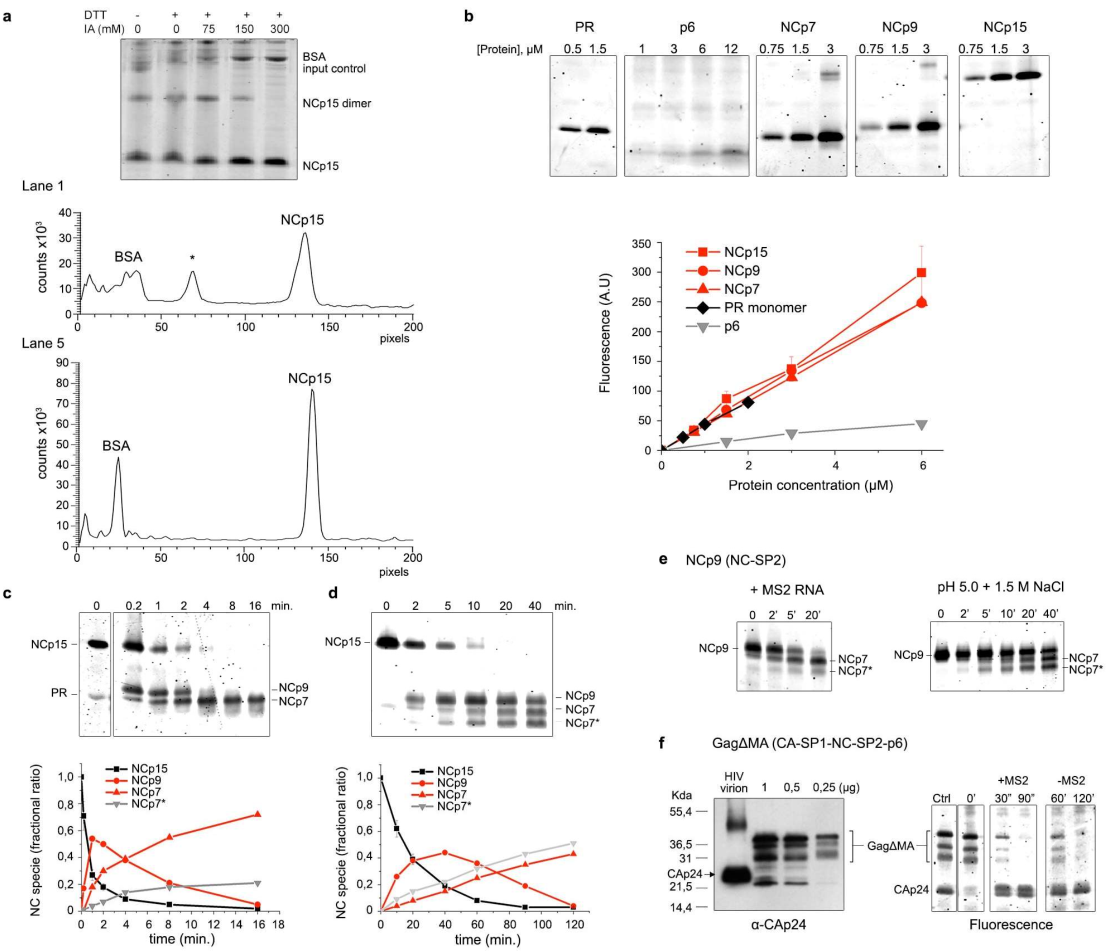


**Supplementary Figure 3 | The PR-driven cleavage of NCp *in vitro* is strongly activated by NA. a**, NC separation by SDS-PAGE and fluorescent staining enables precise quantification of NC species. Boiling in presence of 0.1M DTT and Iodoacetamide (IA) resolved issues of NCp15 oxidization into dimers and increased the fluorescence intensity signal. An example of densitogram is provided between lane 1 and lane 5. **b**, SDS-PAGE separation of PR, p6, NCp7, NCp9 and NCp15. Fluorescence intensity was reported as a function of protein amounts and showed a linear signal enabling accurate protein quantification, augmented by the low sensitivity of p6 to the dye. The basic NCp7 migrated around the position of PR monomer. **c-d**, Kinetics of NCp15 (6µM) cleavage in presence of limiting amounts of PR (0.6 µM) show a strong NA-dependent activation at pH 6.25/0.1 M NaCl (**c**) as compared with enzyme optimum at pH 5.0 / 1.5 M NaCl (**d**). **e**, The NA activation is confirmed for NC-SP2 (NCp9, 6 µM) cleavage in presence of MS2 RNA (120 µM, nt.) at pH 6.25 and 0.1M NaCl when compared to PR activity at pH 5.0 in 1.5 M NaCl. **f,** The SP1-NC cleavage in a GagΔMA construct is also activated by NA. Left panel: Western blot with anti-CAp24 antibodies. The p6-containing GagΔMA protein was very difficult to produce and contained several discrete species. Right panel: products of cleavage (substrate 6µM) for the indicated times in presence or absence of MS2 RNA (120 µM, nt) at pH 6.25 and 0.1M NaCl. Despite substrate heterogeneity, we followed the discrete formation of 25-24 kDa CA-SP1/CA products that clearly revealed SP1-NC cleavage to be strongly activated in the presence of MS2 RNA.

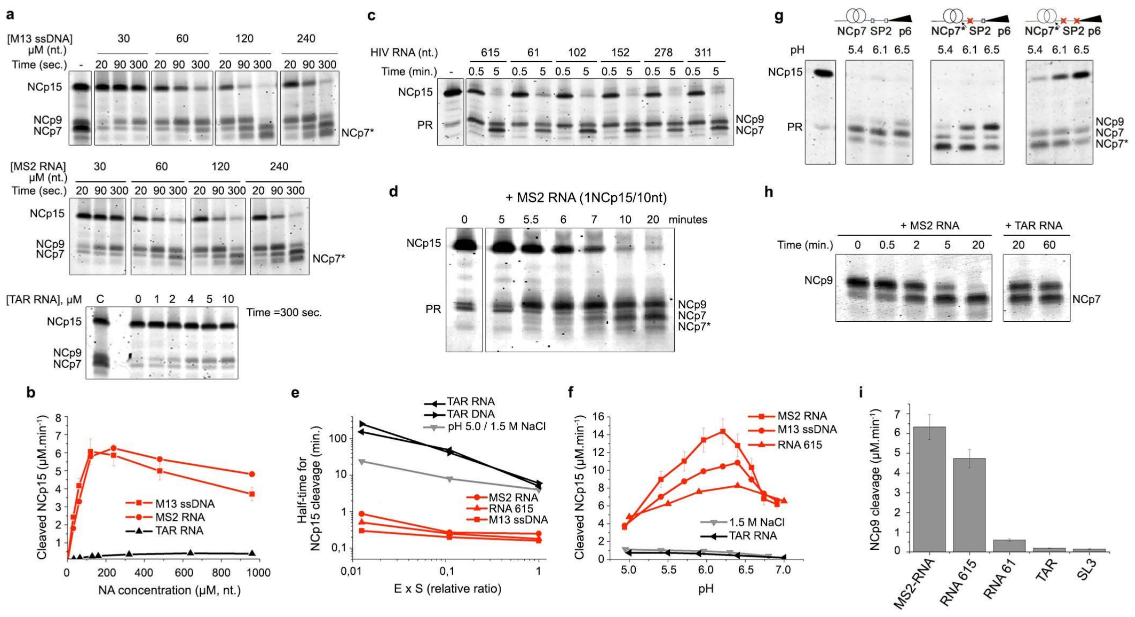


**Supplementary Figure 4 | PR activation of NC proteolysis is modulated by NA length and NC:NA interactions. a-b,** The kinetics of NCp15 cleavage is strongly activated by long ssNA chains (M13 ssDNA and MS2 RNA) but not by short stem-loops like TAR-RNA. In **b**, the apparent reaction rates from the SDS-PAGE analysis in **a** were plotted against NA concentration. **c,** NCp15 cleavage is dependent on RNA-length upon incubation with HIV-1 genomic RNA fragments produced by *in vitro* transcription. **d,** In an environment unfavorable for PR dimer stability (0.1 M NaCl, pH 6.25), NCp15 proteolysis is immediately activated after addition of MS2 RNA to a mixture containing both NCp15 and PR pre-incubated for 5 min. **e,** The crowding effect of the NP complex formed between NCp15 and long ssNA sequesters PR. The half-time for NCp15 cleavage was plotted against the dilution of NCp15 and PR for a fixed NCp15:PR ratio of 10, in presence of M13 ssDNA, MS2 RNA, HIV-1 615-nt RNA, TAR RNA and cTAR DNA. A control reaction was performed at pH 5.0 / 1.5 M NaCl. **f,** The rate of SP2-p6 cleavage was followed as a function of pH and shows a maximum for pH 6.25-6.5 in presence of long ssNA in comparison with TAR RNA or high salts. **g,** An additional cleavage after amino acid 49 (NCp7*) was observed when combining ssNA and low pH, which destabilizes zinc coordination. NCp15 (1) and NCp15 (2) were incubated with PR for 10 min in 0.1M NaCl and at the indicated pH. **h-i**, NCp9 (NC-SP2) cleavage into NCp7 is also activated by a ssNA length-dependent phenomenon involving molecular crowding.

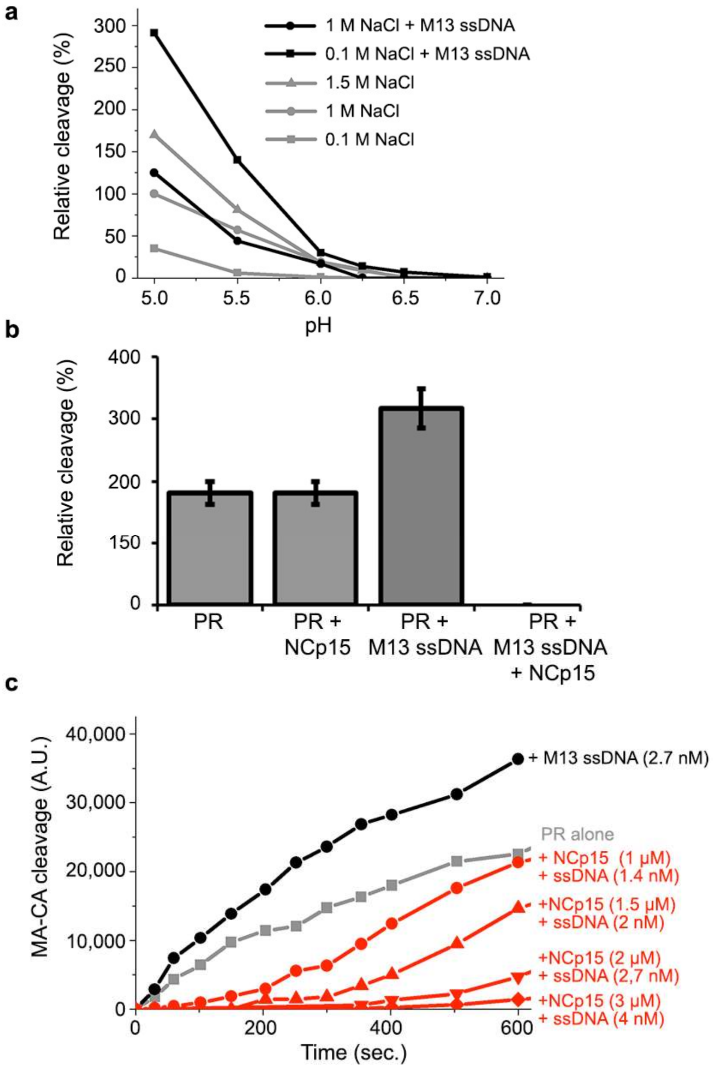


**Supplementary Figure 5 | PR is sequestered by the NCp15:ssDNA NP complex. a,** Followed by FRET, HIV-1 PR is activated by the presence of M13 ssDNA in low salts (0.1M NaCl) , as shown with cleavage of an MA-CA probe as a function of pH. **b,** NCp15 does not compete for the MA-CA probe cleavage at pH 5.5 in 0.1M NaCl, but the NCp15:M13 ssDNA RNP sequesters PR. Cleavage reactions were performed for 5 min. **c,** Increasing the concentration of NCp15:ss M13 NP complex (1 NCp15/10 nt.) sequesters PR, which delays MA-CA cleavage at pH 6.25 in 0.1 M NaCl.

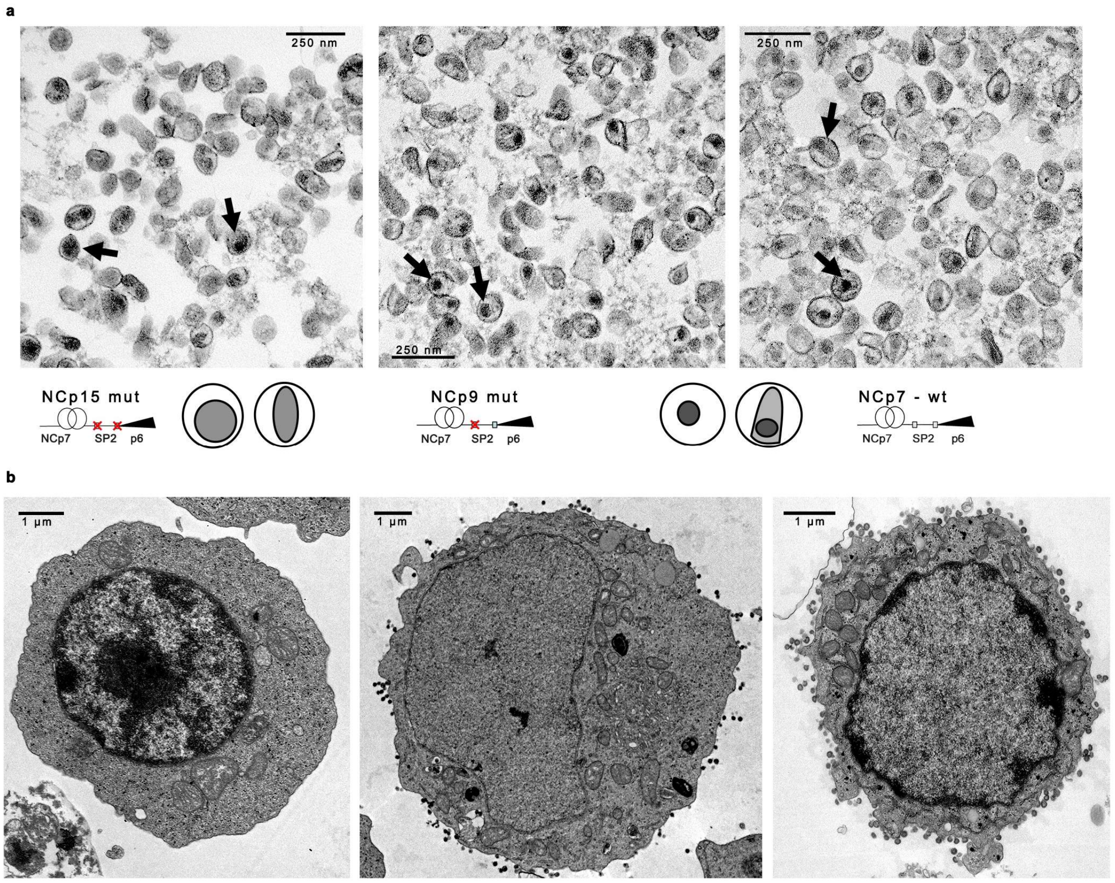


**Supplementary Figure 6 | Nucleocapsid condensation is concomitant with budding. a,** Fields of free HIV-1$_{NL4\text{-}3}$ particles accumulating NCp15, NCp9 or NCp7 (wt virus) show defects in the nucleocapsid condensation for NCp15-containing viruses. The schemes highlight the difference between diffuse cores (examples designed by arrows in NCp15 image) and dense cores (arrows in NCp9 and NCp7). **b,** Multiple washing and fast fixation with glutaraldehyde concentrate virus particles at the plasma membrane of latently infected ACH2 cells (left), producing mature (middle) or immature (right) viruses after 48h activation with vorinostat.

| Parameter | Value | | |
|---|---|---|---|
| | *Case1: PR + MACA* | *Case2: PR + MACA M13* | *Case3: PR + MACA NCp15 + M13* |
| *Initial concentrations and NA size* | | | |
| $n_l$ | - | 7500 | 7500 |
| $[n_t]$ (μM) | - | 20.0 | 20.0 |
| $[S^T{}_1]_0$ ($[MACA^T]_0$) (μM) | 3.0 | 3.0 | 3.0 |
| $[S^T{}_2]_0$ ($[NCp15^T]_0$) (μM) | - | - | 2.0 |
| $[E^T{}_{1/2}]_0$ ($[PR^T{}_{1/2}]_0$) (μM) | 0.1 | 0.1 | 0.1 |
| *Enzyme decay* | | | |
| $K_D$ (nM) | 14.8 | 14.8 | 14.8 |
| $K_R$ (μM) | 2.0 | 2.0 | 2.0 |
| $k^*_{cat}$ ($s^{-1}$) | 0.6 | 0.01 | 0.01 |
| *Enzyme RNP absorption equilibrium* | | | |
| $K^0{}_E$ ($K^0{}_{PR}$) | 0.1 | 0.1 | 0.1 |
| $c_{crit}$ | 3 | 3 | 3 |
| $\xi$ | 1.2 | 1.2 | 1.2 |
| $f$ | 3 | 3 | 3 |
| *Substrate RNP absorption equilibrium* | | | |
| $K_{S1}$ ($K_{MACA}$) | 0.3 | 0.3 | 0.3 |
| $K_{S2}$ ($K_{NCp15}$) | 100 | 100 | 100 |

**Supplementary Table 1 | Kinetic parameters for RNP-modulated two-substrate model.**

### Supplementary Note: A polymer model of nucleic acid-modulated enzyme kinetics

#### 1: Theoretical Development

In order to account for the two phenomena of acceleration and sequestration observed in our *in vitro* biochemical assays (Figure 3 and Supplementary Figure 4), we developed a polymer reaction model of NA-modulated enzyme kinetics. The model consists of a mass-action reaction kinetics approach to follow the competitive processing of two substrates (S1 and S2) by an enzyme (E) in the presence of NA, taking into account equilibrium absorption of each species into the volume pervaded by the NA. It thus enables us to follow the reactions in both the NA pervaded and unpervaded volumes individually whilst relating the transmission of each species across the two domains. We also derive an expression for the relation of the enzyme absorption equilibrium in terms of the experimentally observed phase change in reaction rate upon increased NA length (Figure 3b) in the non-competitive regime. Importantly, the enzyme absorption equilibrium constant is non-linearly dependent on the contiguity of non-specifically bound substrate to individual NA chains. This affords a change in enzyme absorption equilibrium constant as the reaction progresses and is what enables initial uptake of the enzyme by the NA followed by expulsion in the processing-complete regime. We also include the decay of the enzyme due to self-processing. The viral PR is a dimer that can cleave its monomeric units. Therefore, the dimerization of the enzyme is important and described by a characteristic half-life. Including the effects of NA modulation of the enzyme dimer equilibrium improves the accuracy of the model. We fit our one-substrate model to the experimental data to account for the basic feature of length-dependent acceleration and then re-fit parameters to solve the two-substrate model, which directly accounts for sequestration. The predicted reaction kinetics fit the experimental competitive substrate sequestration curves in all three cases (S1 + E, NA + S1 + E, NA + S1 + S2 + E), thus enabling the reaction time for completion of the NA bound substrate reaction to be calculated directly from the model. The model is recalculated for enzyme and substrate conditions characteristic of the *in virio* concentrations to provide an estimate of the timescale of core condensation, which occurs directly upon NCp15 cleavage.

#### 1.1: Pervaded Volume

We first derive an expression for the pervaded volume of a concentration of NA in solution. We assume that we are in a regime of no chain entanglement (due to low concentration and net Coulombic inter-NA repulsion), as exhibited by our AFM experiments (Figure 1). Thus each NA chain can be approximated as occupying an independent volume dependent on its polymer coil properties. Consider a solution of $n_r$ single-stranded NA molecules each composed of $n_l$ nucleotides such that the total nucleotide concentration is $[n_t]$. Thus:

$$n_r = N_A[n_t]V_t/n_l \tag{1}$$

where $V_t$ is the total reaction volume and $N_A$ is Avogadro's constant. We assume that the NA is a flexible Gaussian coil with a persistence length $l_p \sim 2\ l_0 \sim 1.5$ nm, where $l_0 \sim 0.75$ nm is the nucleotide length. The radius of gyration, $R_g$, of a single NA chain is approximated by:

$$R_g \sim (\frac{L.l_p}{3})^{1/2}, \tag{2}$$

where $L = n_l l_0$ is the linear chain length of each NA molecule. Thus the pervaded volume of a single chain $V_r$ can be written as:

$$V_r = \kappa(\frac{n_l l_0 l_p}{3})^{3/2}, \tag{3}$$

and where $\kappa$ is a volumetric constant. The total pervaded volume by all NA, $V_p$, is given by:

$$V_p = n_r V_r. \tag{4}$$

Substituting Equations 1 and 3 into Equation 4 and rearranging, we derive an expression for a dimensionless parameter $\alpha_P$, which corresponds to the ratio of the NA-pervaded volume to the total reaction volume:

$$\alpha_P = V_p/V_t = \kappa N_A [n_t](n_l)^{1/2}(\frac{l_0 l_p}{3})^{3/2}. \tag{5}$$

Correspondingly, the fractional unpervaded volume $\alpha_U$ is simply:

$$\alpha_U = 1 - \alpha_P. \tag{6}$$

Our kinetic model thus consists of a pervaded and unpervaded region whose volume is governed by the polymer properties of the NA (Figure 4a). Although the model consists of multiple pervaded domains within an unpervaded region, these can be summed to represent just one pervaded domain, assuming volumetric homogeneity between individual pervaded volumes.

**1.2: Enzyme-substrate reaction rate equations**

Let us now consider two substrate species ($S_1$ and $S_2$) and one enzyme species ($E$) that are absorbed into and from the pervaded NA volume with equilibrium constants, $K_{S1}$, $K_{S2}$ and $K_E$ , respectively. We thus have the following relations:

$$K_E = [E_P]/[E_U], \tag{7a}$$

$$K_{S1} = [S_{1P}]/[S_{1U}], \tag{7b}$$

$$K_{S2} = [S_{2P}]/[S_{2U}], \tag{7c}$$

where the subscripts $P$ and $U$ refer to the corresponding concentrations of $E$, $S_1$ and $S_2$ in the pervaded (P) and unpervaded domains (U) respectively. We next consider that both $S_1$ and $S_2$ react competitively with $E$ generating intermediate complexes $ES_1$ and $ES_2$ respectively. Given a quasi-steady-state (QSS) approximation, the concentration of each intermediate complex can be expressed in terms of the corresponding Michaelis constants ($K_{M1}$ and $K_{M2}$) for $S_1$ and $S_2$ respectively (see Supplementary Appendix). Then as this intermediate state is short-lived, we can assume that $ES_1$ and $ES_2$ do not cross between domains $U$ and $P$. From this we derive (see Supplementary Appendix) an expression for the total enzyme concentration, $[E_P]$ and $[E_U]$ in domains $P$ and $U$ respectively:

$$[E_P] = [E_T]/[\alpha_P Q_P + \frac{\alpha_U}{K_E} Q_U], \quad (8)$$

$$[E_U] = [E_T]/[\alpha_P K_E Q_P + \alpha_U K_E Q_U], \quad (9)$$

where $[E_T]$ is the absolute total active enzyme concentration and where $Q_P$ and $Q_U$ are the reaction scaling factors in domains $P$ and $U$, owing to the enzyme being distributed across all possible reactions, and are given by:

$$Q_P = 1 + \frac{[S_{1P}]}{K_{M1}} + \frac{[S_{2P}]}{K_{M2}}, \quad (10)$$

$$Q_U = 1 + \frac{[S_{1U}]}{K_{M1}} + \frac{[S_{2U}]}{K_{M2}}. \quad (11)$$

Similarly, approximate relations for the substrate concentrations in the pervaded ($[S_{1P}]$ and $[S_{2P}]$) and unpervaded ($[S_{1U}]$ and $[S_{2U}]$) volumes can be derived, ignoring negligible terms due to the Michaelis complexes with the assumption that $[ES_{1P}] << [S_{1P}]$ and $[ES_{1U}] << [S_{1U}]$ (see Supplementary Appendix):

$$[S_{1P}] \sim [S_{1T}]/(\alpha_P + \frac{\alpha_U}{K_{S1}}), \quad (12)$$

$$[S_{1U}] \sim [S_{1T}]/(\alpha_P K_{S1} + \alpha_U), \quad (13)$$

$$[S_{2P}] \sim [S_{2T}]/(\alpha_P + \frac{\alpha_U}{K_{S2}}), \quad (14)$$

$$[S_{2U}] \sim [S_{2T}]/(\alpha_P K_{S2} + \alpha_U), \quad (15)$$

where $[S_{1T}]$ and $[S_{2T}]$ are the total substrate concentrations of each species. We then follow the four rate equations, one for each of the two species in each of the two domains in terms of the corresponding concentrations within each domain:

$$\nu_{1P} = -\frac{d[S_{1P}]}{dt} = -\frac{k_{cat1}}{K_{M1}}[E_P][S_{1P}], \quad (16)$$

$$\nu_{1U} = -\frac{d[S_{1U}]}{dt} = -\frac{k_{cat1}}{K_{M1}}[E_U][S_{1U}], \tag{17}$$

$$\nu_{2P} = -\frac{d[S_{2P}]}{dt} = -\frac{k_{cat2}}{K_{M2}}[E_P][S_{2P}], \tag{18}$$

$$\nu_{2U} = -\frac{d[S_{2U}]}{dt} = -\frac{k_{cat2}}{K_{M2}}[E_U][S_{2U}], \tag{19}$$

where $k_{cat1}$ and $k_{cat2}$ are the and turnover numbers for $S_1$ and $S_2$ respectively.

#### 1.3: Enzyme decay

Viral PR decays with a well-characterized half-life of 30 mins[34]. This is due to recognition and inactivating cleavage of PR monomers by the active dimer. Therefore the dimerization equilibrium of PR is expected to play a role in concentration of active PR available to be sequestered into the NA pervaded volume. Generalizing to enzyme $E$, consisting of a dimer of two monomers $E_{1/2}$, we have the following reaction schemes:

$$E_{\frac{1}{2}} + E_{\frac{1}{2}} \overset{K_D}{\rightleftharpoons} E, \tag{20}$$

$$E + E_{\frac{1}{2}} \overset{K_R}{\rightleftharpoons} E.E_{\frac{1}{2}} \overset{k^*_{cat}}{\rightarrow} E + E^*_{\frac{1}{2}}, \tag{21}$$

where $K_D$ is the enzyme dimer dissociation constant and where $K_R$ and $k^*_{cat}$ are the Michaelis constant and turnover numbers of the enzyme inactivation reaction, $E.E_{1/2}$ is the steady-state intermediate and $E^*_{1/2}$ is the cleaved monomer. We then have the following relations between each species:

$$[E] = \frac{[E_{\frac{1}{2}}]^2}{K_D}, \tag{22}$$

$$[E.E_{\frac{1}{2}}] = \frac{[E_{\frac{1}{2}}]^3}{K_D K_R}. \tag{23}$$

The total monomer concentration ($[E^T_{1/2}]$) is given by:

$$[E^T_{\frac{1}{2}}] = \frac{3[E_{\frac{1}{2}}]^3}{K_D K_R} + \frac{2[E_{\frac{1}{2}}]^2}{K_D} + [E_{\frac{1}{2}}], \tag{24}$$

and the decay of viable monomers is then determined by the following rate law:

$$\frac{d[E_{\frac{1}{2}}]}{dt} = -\frac{k^*_{cat}[E_{\frac{1}{2}}]^3}{K_D K_R}. \quad (25)$$

**1.4: Dependence of enzyme absorption equilibrium constant on RNP contiguity**

We define the RNP contiguity number ($c$) as the average number of non-specifically bound substrate molecules within the pervaded volume ($S_{2P}$) per NA chain, consisting of $n_l$ nucleotides. Then for $K_{S2} >> 1$, the overwhelming majority of $S_2$ are in the pervaded volume and the contiguity can be expressed as:

$$c \sim \frac{[S_{2T}]n_l}{[n_t]}, \quad (26)$$

where $[S_{2T}]/[n_t]$ is the ratio of total RNP binding substrate concentration to total nucleotide concentration. Consider enzyme $E$ absorbing into the RNP pervaded volume with absorption rate constant $k_e$ and escaping with a rate constant $k_{-e}$ to yield equilibrium constant $K_E = k_e / k_{-e}$. We stipulate that increases in contiguity result in an increased mean time for enzyme escape, thus a decrease in the escape constant $k_{-e}$, but not the absorption rate $k_e$. Then for NA with little or no bound substrate, that corresponds to a contiguity less than the critical threshold $c_{crit}$ ($c < c_{crit}$), we have an invariant baseline escape constant $k_{-e} = k^0_{-e}$ and thus equilibrium constant $K^0_E = k_e / k^0_e$. Beyond the critical threshold ($c > c_{crit}$), $k_{-e}$ becomes non-linearly dependent on contiguity $c$, with exponent ξ and constant $f$ such that:

$$K_E = K^0_E(f + \frac{c}{c_{crit}})^{\xi}. \quad (27)$$

Thus in our model, enzyme absorption is explicitly dependent on RNP-bound substrate number (contiguity) - then as this substrate is cleaved over time, we expect a dramatic decrease in the RNP's capacity to absorb the enzyme and this directly affects the enzyme reaction rate equations.

**2: Computational implementation and model parameters**

The theoretical model was implemented computationally in MATLAB. The coupled rate equations were numerically solved using the fourth-order Runge-Kutta ordinary differential equation (ODE) solver in MATLAB with a timestep of 1 second. Each integration timestep was split into two sequential components - first to compute the instantaneous active enzyme dimer concentration using the decay equation (Eq. 25) and secondly, this concentration was input into the enzyme-substrate reaction equations to compute the update on substrate concentrations.

**3: Analysis of model features**

The default general enzyme reaction parameters used in this study were $k_{cat1}=k_{cat(MACA)}$ = 7.07 $s^{-1}$, $k_{cat1}=k_{cat(MACA)}$ = 0.65 $s^{-1}$, $K_{M1}=K_{M(MACA)}$= 0.14 mM and $K_{M2}=K_{M(NCp15)}$ = 0.03 mM. The polymer model parameters were $l_p$ =75.0 nm, $l_0$=1.0 nm and κ = 0.1.

#### 3.1: Sensitivity of pervaded volume to physical polymer properties

The variation of the pervaded volume with varying nucleotide chain length $n_l$ was calculated for default polymer coil properties at a range of values for total nucleotide concentration $[n_t]$ in the approximate range (10 – 200 μM) carried out in the experimental assays. The effect of varying persistence length was also determined.

For a given nucleotide concentration, increasing chain length increases pervaded volume according to the $(n_l)^{1/2}$ dependency. However, in the relevant range of concentrations considered experimentally (10 – 200 μM), $\alpha_P$ does not increase beyond 0.1 even for NA approaching 1000 nucleotides. At 20 μM, $\alpha_P$ is below 0.02 for even 10,000 nucleotide chains. Increasing persistence length $l_p$ by 2-fold and 10-fold for $[n_t]$ =10 μM, thus relaxing the polymer coiling propensity, still does not increase $\alpha_P$ beyond 0.02 and 0.1 for 10,000 and 1,000 nucleotide chain lengths respectively. Therefore, according to our polymer model at moderate nucleotide concentration, the pervaded volume is always a small fraction of the overall reaction volume.

#### 3.2: Matching a one-substrate model to experimental rate observables

We developed a one-substrate model to compare against the corresponding experimental assay at varying nucleotide length, $n_l$ (Figure 4b). Nucleotide concentration was set to $[n_t]$ =120 μM, initial total NCp15 concentration was $[S^T{}_2]_0$=$[NCp15^T]_0$=6 μM, initial total PR monomer structure was $[E^T{}_{1/2}]_0$= $[PR^T{}_{1/2}]_0$=1.2 μM. Enzyme decay parameters were set to: $K_D$=14.8 nM, $K_R$=2.0 μM and $k^*_{cat}$ =0.01 $s^{-1}$. The enzyme RNP absorption equilibrium parameters were set to: $K^0{}_E$=$K^0{}_{PR}$=0.54. The NCp15 RNP absorption equilibrium parameter was set to $K_{S2}$=$K_{NCp15}$=100.

Typically, the change of substrate concentrations in each of the separate reaction volumes (pervaded and unpervaded) cannot be followed directly by experiment. Therefore, to establish a direct comparison with experimental observables, it is necessary to reformulate Equations 16-19 in terms of the change in total substrate concentration $v_{1T}$ and $v_{2T}$. In a one-substrate model consisting of just $S_2$, in this case corresponding to NCp15, we recast Eq 18 in terms of Eqs 8 and 14 and rearranging we can write the total reaction rate of processing $S_2$ as:

$$\nu_{2T} = \frac{1}{A + (B/K_E)} \tag{28}$$

Where $A$= $\gamma\alpha_P Q_P$ and $B$ = $\gamma\alpha_U Q_U$ and where $\gamma$ = $K_{M2}/(k_{cat2}\ [E_T]\ [S_{2T}])$. We further express $A$ and $B$ in terms of $\alpha_P$ and the total substrate concentration $[S_{2T}]$ and the total enzyme concentration $[E_T]$:

$$A = \gamma\alpha_P\Big(1 + \frac{[S_{2T}]K_{S2}}{(1+(K_{S2}-1)\alpha_P)K_{M2}}\Big) \tag{29}$$

$$B = \gamma\alpha_P\Big(1 + \frac{[S_{2T}]}{(1+(K_{S2}-1)\alpha_P)K_{M2}}\Big). \tag{30}$$

Note that this gives:

$$A + B = \frac{K_{M2}}{k_{cat2}}[E_T][S_{2T}]\Big(1 + \frac{[S_{2T}]}{K_{M2}}\Big), \tag{31}$$

so that in the case that $K_E$ = 1 we recapture the normal Michaelis-Menten equation from Equation 28. This formulation enables a direct handle on experimentally observable changes in total substrate concentration [$S_{2T}$] in terms of the substrate equilibrium absorption properties, $K_{S2}$, as well as those of the enzyme, $K_E$. It also permits prediction of reaction rate in terms of nucleotide chain length-dependence, $n_l$, which controls $\alpha_P$.

#### 3.2.1: Estimation of initial enzyme absorption equilibrium

In the specific case that we are in the sub-critical contiguity regime ($c < c_{crit}$), then $K_E = K^0_E$ and if we assume $v = v_0$ = 0.02 mM/s corresponding to the $n_l$-independent regime of the experimental assay as shown in Figure 4b, then $K^0_E = B/(1/v_0 - A)$. Although $K^0_E$ is formally dependent on $\alpha_P$ and thus $n_l$, it is extremely insensitive to variation of $\alpha_P$. Therefore, even order of magnitude changes in NA persistence length and the volumetric constant do not affect $K^0_E$, which can effectively be treated as a constant. For $n_l$ in the $n_l < n_{l,crit}$ regime we obtain mean $K^0_E$ = 0.545 +/- 0.005, where we take $n_{l,crit} \sim 50$ as estimated from experiment. This provides a fitted estimate for $K^0_E$ in terms of the experimental data.

#### 3.2.3: Fitting of acceleration assay to one substrate model

Using the experimentally fitted value for $K^0_E$ and $n_{l,crit}$ , which gives $c_{crit} \sim 1.1$, for the low-$n_l$ constant rate ($v_0$) regime, we fit the $n_l$-dependent experimental data to Eq 28 using the general definition of $K_E$ in terms of Eq 27, using parameters $f$ and $\xi$ and obtain $f = 3$, $\xi = 0.4$ (Figure 4b). Our one-substrate model is therefore consistent with the notion that beyond a critical threshold the enzyme absorption equilibrium becomes non-linearly dependent on the contiguity of substrate molecules within the NA pervaded volume resulting in the acceleration of substrate processing, whilst below it a constant reaction rate is exhibited.

#### 3.3: Two-substrate model

Given that the basic feature of NA length dependent acceleration is already captured in the one-substrate model, we next evaluated whether the same theoretical approach could account for the sequestration effect that was observed experimentally using an additional competitive substrate. Here, rather than fitting reaction rates at half-original concentrations, we require computing the time evolution of substrate concentration of both substrates whilst also fitting the required parameters of our model. The fitted parameters used for the two-substrate model reactions are provided in Supplementary Table 1.

### Supplementary Appendix

Consider a total reaction volume $V_t$ partitioned into two effective sub-volumes $V_p$ and $V_u$ in each of which occur competitive enzymatic reactions between one enzyme $E$ and two substrate species $S_1$ and $S_2$. This constitutes four reactions described by the following scheme:

Reactions in sub-volume $V_p$:

$$E_P + S_{1P} \underset{k_{-1,1}}{\overset{k_{1,1}}{\rightleftharpoons}} ES_{1P} \overset{k_{cat1}}{\rightarrow} E_P + P_{1P},$$

(A1)

$$E_P + S_{2P} \underset{k_{-1,2}}{\overset{k_{1,2}}{\rightleftharpoons}} ES_{2P} \overset{k_{cat2}}{\rightarrow} E_P + P_{2P}, \tag{A2}$$

Reactions in sub-volume $V_u$:

$$E_U + S_{1U} \underset{k_{-1,1}}{\overset{k_{1,1}}{\rightleftharpoons}} ES_{1U} \overset{k_{cat1}}{\rightarrow} E_U + P_{1U}, \tag{A3}$$

$$E_U + S_{2U} \underset{k_{-1,2}}{\overset{k_{1,2}}{\rightleftharpoons}} ES_{2U} \overset{k_{cat2}}{\rightarrow} E_U + P_{2U}, \tag{A4}$$

where subscripts P and U denote the corresponding species in each of the separate sub-volumes $V_p$ and $V_u$ respectively, $ES_1$ and $ES_2$ are the Michaelis complexes and $P_1$ and $P_2$ are the products. The total enzyme number $E^T$ is then:

$$E^T = E^P + ES_1^P + ES_2^P + E^U + ES_1^U + ES_2^U \tag{A5}$$

where superscripts P and U denote the absolute number of each species. Dividing by the total volume we obtain:

$$\frac{E^T}{V_t} = \alpha_P \frac{E^P}{V_P} + \alpha_P \frac{ES_1^P}{V_P} + \alpha_P \frac{ES_2^P}{V_P} + \alpha_U \frac{E^U}{V_U} + \alpha_U \frac{ES_1^U}{V_U} + \alpha_U \frac{ES_2^U}{V_U} \tag{A6}$$

where $\alpha_P = V_p/V_t$ and $\alpha_U = V_U/V_t$ are the fractional sub-volumes for P and U respectively. Then we use subscripts P and U to represent the effective concentration of each species in its corresponding sub-volume (e.g. $[E_T] = E^T/V_t$, $[E_P] = E^P/V_p$, $[ES_{1P}] = ES_1^P/V_p$, etc.) and by factoring we obtain the total enzyme concentration $[E_T]$ in terms of the effective concentrations in each sub-volume:

$$[E_T] = \alpha_P([E_P] + [ES_{1P}] + [ES_{2P}]) + \alpha_U([E_U] + [ES_{1U}] + [ES_{2U}]) \tag{A7}$$

Assuming a quasi-steady state for each reaction in each of the sub-volumes and that the reaction transition is sufficiently short-lived that $ES_1$ and $ES_2$ do not cross between domain U and P, we can rewrite $[ES_{1P}]$, $[ES_{2P}]$, $[ES_{1U}]$ and $[ES_{2U}]$ in terms of the respective Michaelis constants ($[ES_{1P}] = [E_P][S_{1P}]/K_{M1}$, $[ES_{2P}] = [E_P][S_{2P}]/K_{M2}$, $[ES_{1U}] = [E_U][S_{1U}]/K_{M1}$ and $[ES_{2U}] = [E_U][S_{2U}]/K_{M2}$). Substituting these into the above equation and rearranging we have:

$$[E_T] = \alpha_P[E_P]\Big(1 + \frac{[S_{1P}]}{K_{M1}} + \frac{[S_{2P}]}{K_{M2}}\Big) + \alpha_U[E_U]\Big(1 + \frac{[S_{1U}]}{K_{M1}} + \frac{[S_{2U}]}{K_{M2}}\Big) \tag{A8}$$

Then by identifying the terms in brackets as $Q_P$ and $Q_U$ respectively as expressed in equations 10 and 11 and substituting out either [$E_P$] or [$E_U$] using the enzyme equilibrium equation (7a) we obtain Equations 8 and 9 for the effective enzyme concentrations in each sub-volume.

Using a similar approach to above, the total substrate number, $S_1^{\mathrm{T}}$, for $S_1$ is:

$$S_1^T = S_1^P + ES_1^P + S_1^U + ES_1^U, \tag{A9}$$

from which we can express the total concentration [$S_{1T}$] as:

$$[S_{1T}] = \alpha_P([S_{1P}] + [ES_{1P}]) + \alpha_U([S_{1U}] + [ES_{1U}]). \tag{A10}$$

By rewriting [$ES_{1P}$] and [$ES_{1U}$] in terms of the Michaelis constants and by making use of the substrate equilibrium equation (7b) to substitute out [$S_{1U}$], and rearranging we obtain:

$$[S_{1T}] = [S_{1P}]\Big(\alpha_P + \frac{\alpha_U}{K_{S1}}\Big) + [S_{1P}]\Big(\frac{\alpha_P[E_P]}{K_{M1}} + \frac{\alpha_U[E_U]}{K_{M1}K_{S1}}\Big). \tag{A11}$$

By further substituting out [$E_U$] using the enzyme equilibrium equation (7a) we obtain:

$$[S_{1T}] = [S_{1P}]\Big(\alpha_P + \frac{\alpha_U}{K_{S1}}\Big) + \frac{[E_P][S_{1P}]}{K_{M1}}\Big(\alpha_P + \frac{\alpha_U}{K_E K_{S1}}\Big). \tag{A12}$$

This yields an exact expression for [$S_{1P}$] in terms of the total substrate concentration [$S_{1T}$]:

$$[S_{1P}] = [S_{1T}]/\Big(\alpha_P + \frac{\alpha_U}{K_{S1}} + \frac{[E_P]}{K_{M1}}(\alpha_P + \frac{\alpha_U}{K_E K_{S1}})\Big). \tag{A13}$$

Similarly, if one substitutes out [$S_{1P}$] in terms of [$S_{1U}$] and [$E_P$] in terms of [$E_U$] from equation A10, one derives [$S_{1U}$]:

$$[S_{1U}] = [S_{1U}]/\Big(\alpha_P K_{S1} + \alpha_U + \frac{[E_U]}{K_{M1}}(\alpha_P K_E K_{S1} + \alpha_U)\Big). \tag{A14}$$

Correspondingly for [$S_{2P}$] and [$S_{2U}$] these are:

$$[S_{2P}] = [S_{2T}]/\Big(\alpha_P + \frac{\alpha_U}{K_{S2}} + \frac{[E_P]}{K_{M2}}(\alpha_P + \frac{\alpha_U}{K_E K_{S2}})\Big). \tag{A15}$$

$$[S_{2U}] = [S_{2U}]/\Big(\alpha_P K_{S2} + \alpha_U + \frac{[E_U]}{K_{M2}}(\alpha_P K_E K_{S2} + \alpha_U)\Big). \tag{A16}$$

Finally, if we assume that [$S_{1P}$] >> [E$S_{1P}$], then equation A12 can be approximated to equation 12 and similarly for [$S_{1U}$], [$S_{2P}$] and [$S_{2U}$] this approximation leads directly to equations 13-15.
